# A hydrocarbon pheromone mediates ant group cohesion

**DOI:** 10.64898/2026.11.25.769233

**Authors:** Matteo Rossi, Tiphaine P. M. Bailly, Jocelyn G. Millar, Dennis O. Melendez, Stephany Valdés-Rodríguez, Thomas Schmitt, Erik T. Frank, Daniel J. C. Kronauer

## Abstract

Social insect colonies are among the most cohesive groups in the animal kingdom, yet we know little about the signals and sensory mechanisms that underlie their collective unity. Here, we show that cuticular hydrocarbons (CHCs), low-volatility compounds present on the insect body surface, play a central role in social cohesion in the clonal raider ant, *Ooceraea biroi*. CHCs are transferred from the body to the nest substrate, where they are sufficient to induce aggregation. Nest-associated CHCs elicit aggregation across distinct *O. biroi* genotypes, indicating that they function as a species-wide signal. In contrast, hydrocarbon mixtures from other ant species or synthetic hydrocarbons absent from the *O. biroi* cuticle do not elicit aggregation. We further show that behavioral responses to nest-associated CHCs vary with age and physiological state and correlate with the ants’ propensity to leave the nest. This suggests that variation in responses to CHCs contributes to age-based division of labor and other behavioral dynamics in the colony. Finally, we show that aggregation is specifically mediated by 11,15-dimethylheptacosane, the most abundant component of the *O. biroi* CHC profile. Together, these findings identify a pheromone-based mechanism underlying group formation and persistence, as well as division of labor in the clonal raider ant.

## Introduction

Solitary animals typically associate only transiently, most often for mating, whereas social animals have evolved mechanisms that enable stable group living. The transition from solitary to social life therefore likely requires profound changes in how individuals process and respond to sensory information. Consistent with this idea, social species are highly attuned to social information, including signals that convey dominance relationships or group affiliation^1,2^. Environmental perturbations can likewise reveal contrasts between solitary and social systems: in *Drosophila* groups, sensing elevated temperatures drives individual escape responses^3^, whereas in ants, similar perturbations trigger colony-level reactions, in which workers respond in a spatiotemporally coordinated manner to evacuate the nest^4^. Chemical communication plays a central role in coordinating ant social life, and the odor signals (pheromones) regulating behaviors such as alarm responses and trail following have been characterized in many species^5,6^. However, although group formation and maintenance are essential to ant fitness^7^, the sensory cues that ants use to establish a nest and that allow them to maintain colony cohesion remain unknown.

Aggregation pheromones, odorants that attract individuals of both sexes and promote group formation, have been identified in diverse invertebrate taxa, including bark beetles, bed bugs, locusts, termites and fruit flies^8,9,10,11,12,13,14^. Surprisingly, no such aggregation pheromone has been described in social hymenopterans, despite their heavy reliance on olfaction for social coordination. Nonetheless, evidence suggests that olfactory cues are essential for ant group formation: in both the clonal raider ant, *Ooceraea biroi*, and the Indian jumping ant, *Harpegnathos saltator*, two distantly related species, workers carrying mutations that disrupt odorant receptor function no longer join colonies^15,16^. This implicates odor detection as essential for ant social cohesion.

Here, we study the chemical ecology of aggregation and nest formation in *O. biroi*, an experimentally tractable model system^17,18^. The species is queenless, and workers reproduce clonally^19,20^. Colonies of *O. biroi* are subterranean and do not excavate complex nest structures like many other ants. Instead, they form simple nests by densely aggregating in small soil pockets^19,21^. This behavior is recapitulated under laboratory conditions, where groups of workers form dense aggregates around the brood from which they forage and to which they return with food. Colonies undergo phasic cycles in which they alternate between foraging phases, during which larvae stimulate foraging activity in the adults, and reproductive phases, where larvae are initially absent, the colonies contain pupae, and the ants remain inside the nest and lay eggs^21,22^. Like other social insects, *O. biroi* exhibits age-related division of labor, and during the foraging phase, young workers tend to remain inside the nest and nurse the brood while older workers tend to leave the nest and forage^23,24,25^. The tendency of the ants to remain aggregated inside the nest therefore changes both as a function of cycle phase and age.

## Results

### Ants mark the nest with a cue that promotes aggregation

To investigate how ants form and maintain aggregates, we developed a behavioral assay in which adult ants and pupae were placed into featureless arenas with white filter paper surfaces that lacked predefined nest structures. Upon release into the arena, the ants form a dense and spatially stable nest aggregate within minutes **(Fig. 1A)**. We identified and tracked the location of the ants’ nest based on the density of dark pixels (the ants are dark while the background is light) using a custom automated video analysis pipeline. We defined the “nest” as the region containing the highest density of dark pixels **(Fig. 1B)**. We then used this assay to test the hypothesis that the ants rely on sensory cues associated with the nest environment to form and maintain aggregates. A colony composed of 100 old ants in the reproductive phase, together with 50 pupae, was allowed to establish a nest overnight in the arena, after which the ants were removed. The arena was then rotated by 90° to control for the possible influence of cues outside of the arena on nest site choice. A naïve colony of identical composition was then introduced into the same arena **(Fig. 1C)**. Colonies preferentially nested in locations previously occupied by other colonies (Wilcoxon rank-sum test, *P* < 0.001) **(Fig. 1D)**, suggesting that ants leave a cue behind that marks the nest site. This effect became weaker as the interval between removal of the first colony and introduction of the second colony increased, but it remained significant even with an interval of 24 hours (*P* < 0.01) **(Fig. 1D)**. This is consistent with a low-volatility chemical cue that persists for at least 24 hours and guides nest formation.

**Figure 1.**
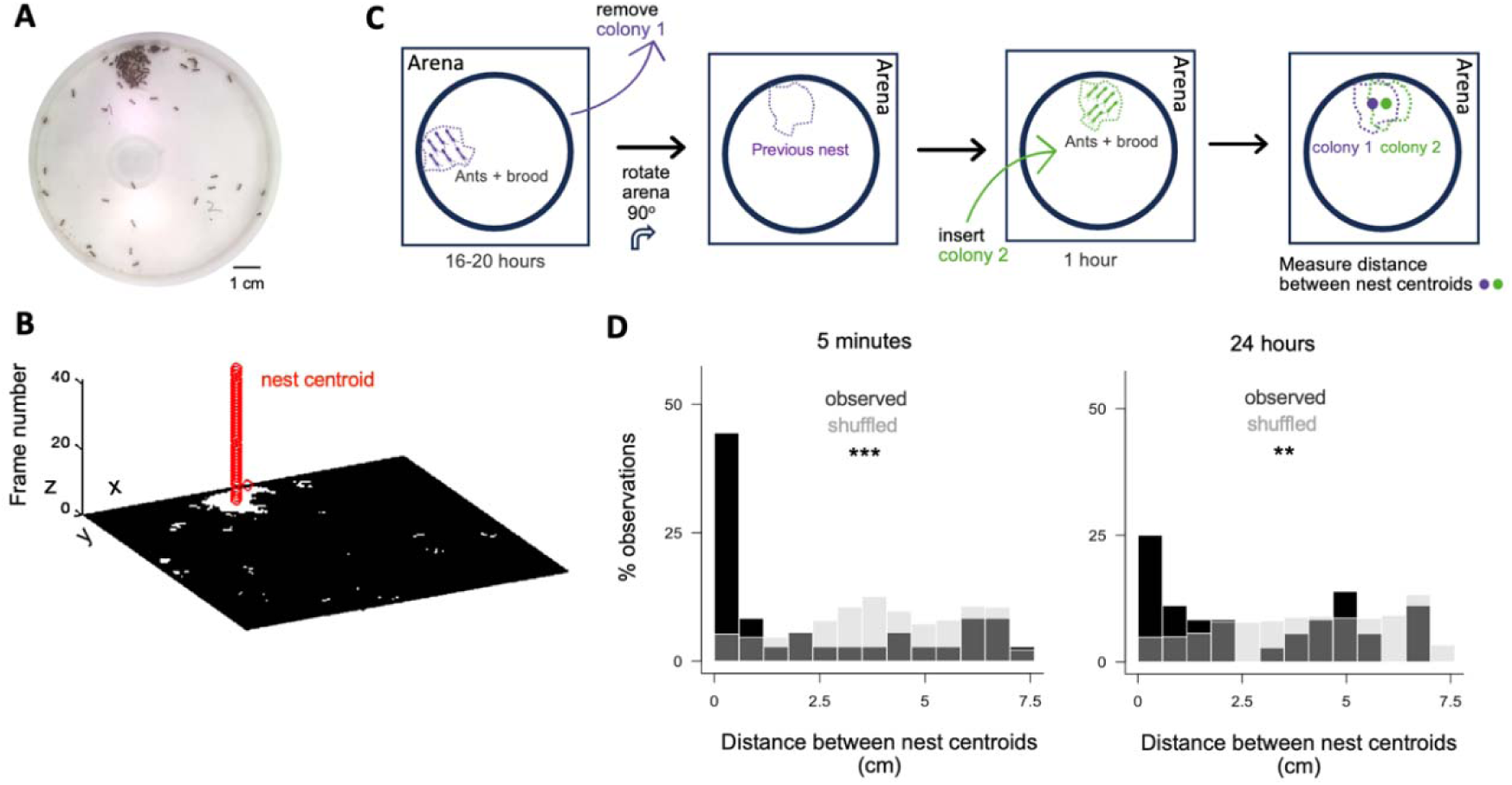
Ants mark the nest with a cue that promotes aggregation. **(A)** Example of the experimental arena (7.6 cm diameter) with an ant colony forming a nest. **(B)** Schematic output of the automated video analysis pipeline used to identify the nest centroid over time, based on the spatial density of ant-colored pixels throughout an hour-long assay. In this rendition, ants are shown in white, the background in black, and the nest centroids are indicated by red circles, with one frame analyzed every 90 seconds. **(C)** Experimental design to test whether colonies reestablish nests at previous nest sites. Colony 1 (purple) is allowed to settle for 16–20 hours before being removed from the arena. After rotating the arena by 90°, colony 2 (green) is introduced and given 1 hour to form a nest. The distance between nest centroids is then measured. **(D)** Distribution of distances between successive nest centroids compared with shuffled controls (null expectations were generated by shuffling centroid positions across independent experiments). The time elapsed between removing colony 1 and adding colony 2 is indicated at the top of each panel. Colonies established nests significantly closer to previous sites than expected by chance, both 5-minutes and 24-hours after colony 1 removal. **, *P* < 0.01. ***, *P* < 0.001. N = 36 replicates for each condition.

To test whether brood is required for the accumulation of nest cues, we repeated the nest-site overlap assay in the absence of pupae. Nest-site overlap remained significantly greater than expected by chance using colonies composed only of 100 old adults in the reproductive phase, indicating that adults alone are sufficient for these cues to accumulate on the nest substrate (**Fig. S1**).

### Behavioral responses to nest cues change with age and phase of the colony cycle

In *O. biroi*, the tendency of ants to leave the nest changes both with age (on average, older ants leave the nest more than younger ants) and phase of the colony cycle (ants leave the nest often during the foraging phase, but rarely during the reproductive phase). To test whether these dynamics could be in part explained by differences in the ants’ responses to nest cues, we quantified group cohesion in experimental arenas using ants of different ages and from different phases of the colony cycle **(Fig. 2A)**. We defined and measured group cohesion as the number of dark pixels located within the nest region divided by the total number of dark pixels in the arena, considering the last 10 frames (1 frame per 30 seconds) of a 3-hour assay (see Methods) **(Fig. 2B)**. Colony cohesion was highest in groups of young ants in the foraging phase, intermediate in older ants in the reproductive phase, and lowest in older ants in the foraging phase (all tested in the absence of brood) (Kruskal-Wallis test *P* < 0.001, pairwise Wilcoxon rank-sum tests with Bonferroni-Holm correction, all *adj. P* < 0.001) **(Fig. 2C)**. These results suggest that the same mechanisms that underlie differences in the ants’ tendency to aggregate in our experimental setup might also determine their tendency to remain inside the nest or to leave the nest to explore and forage in more natural settings.

**Figure 2.**
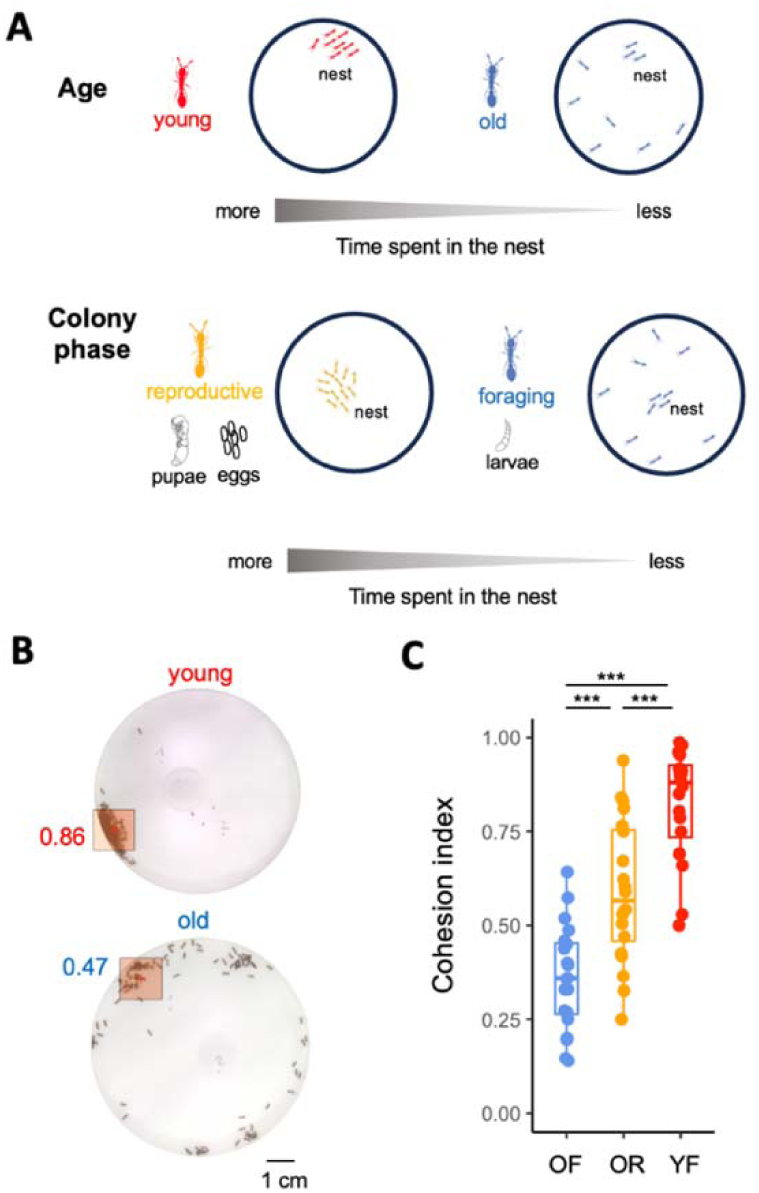
Behavioral responses to nest cues change with age and phase of the colony cycle. **(A)** Schematic summarizing behavioral differences associated with worker age and phase of the colony cycle. On average, young workers spend more time inside the nest than old workers during the foraging phase (top). Ants do not normally leave the nest during the reproductive phase, but they explore and forage outside the nest during the foraging phase (bottom). **(B)** Representative nest images illustrate denser clustering among young (3-7 days old) compared with old (>1.2 months old) workers in the foraging phase. The red squares represent the tiles with the highest density of dark pixels in the arena, the center of which is the nest centroid (see Methods). **(C)** Boxplots of cohesion index across colony types without brood (each colony was composed of 100 ants). Each dot represents one colony composed of old ants in the foraging phase (OF, blue; >1.2 months old), old ants in the reproductive phase (OR, orange; >1.7 months old), and young ants in the foraging phase (YF, red; 3-7 days old) (n = 20 replicates per condition). Boxplots show upper and lower quartiles, whiskers represent 1.5 times the interquartile range, and horizontal bars indicate medians. ***, *adj. P* < 0.001.

### Cuticular hydrocarbons transferred to the nest substrate act as arrestants

To investigate the chemical nature of the aggregation cue, we housed adult ants together with pupae in arenas lined with filter paper. After 24 hours, we removed ants and pupae and cut out the part of the filter paper on which the ants had nested. As a control, we cut out an area of similar size that the ants had had access to, but on which they had not nested **(Fig. S2A)**. Filter paper cutouts were then soaked for 2 hours in one of several solvents spanning a range of polarities and affinities for different classes of chemical compounds **(Fig. S2B)**. We applied the resulting nest and control extracts to a fresh arena, introduced a different ant colony, and monitored its site preference. In each independent behavioral assay, we scored whether the colony aggregated on the nest extract, the control extract, or neither, and calculated the frequency of each response **(Fig. S2B)**. Among the tested solvents, nest extracts with hexane, which is highly non-polar, showed the strongest tendency to induce aggregation compared to the respective control (χ² test, *P* < 0.05), while nest extracts obtained with more polar solvents elicited weaker responses (**Fig. S2B**). These results suggest that nest extracts with hexane contain compounds that induce aggregation.

Gas chromatography-mass spectrometry (GC-MS) of the hexane extracts from nest areas revealed a chemical profile dominated by hydrocarbons that matched the cuticular hydrocarbon (CHC) profile of ant whole body extracts **(Fig. 3A**). This was true for nests composed of adults and brood **(Fig. 3A, Fig. S3)**, and for nests composed only of young or old adults (**Fig. S4A-D**). CHCs are long-chain hydrocarbons coating the insect cuticle. These compounds originally served to prevent desiccation, but they have been co-opted for diverse social signaling roles^26,27^. GC-MS analysis of hexane extracts of *O. biroi* adults, pupae, and larvae revealed the same set of CHCs across developmental stages (with the exception of one compound present only in larvae, **Dataset S1**), with differences in the relative abundance of some compounds, especially between larvae and adults (**Fig. S5A**). We found no significant differences in CHC profiles between old ants in the reproductive and foraging phase (PERMANOVA test, *P* > 0.05) **(Fig. S5B**; see also^28^). These results suggest that the ants’ CHCs get transferred from the body surface to the substrate, where they persist and serve as arrestant nest cues. To test this hypothesis directly, we immersed adult ants in hexane for ten minutes to extract nonpolar cuticular chemicals and applied these extracts to our behavioral arenas (**Fig. 3B**). Cuticular extracts, composed primarily of CHCs, had strong arrestant effects and induced aggregation (χ² test, *P* < 0.01) **(Fig. 3C**).

**Figure 3.**
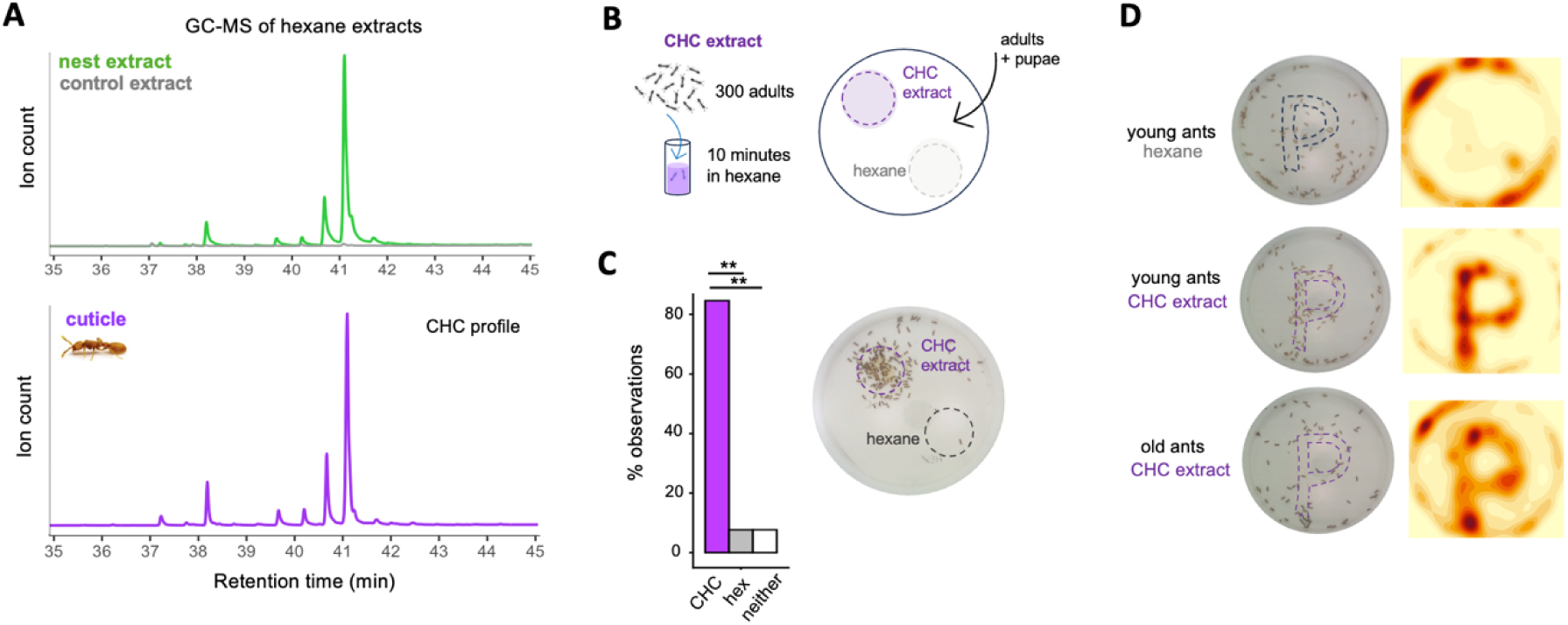
Cuticular hydrocarbons transferred to the nest induce arrestment in ants. **(A)** GC–MS profiles of hexane extracts from nest (green) and control (gray) patches of filter paper (top), and of a hexane surface extract from an *O. biroi* worker (purple, bottom). The chemical profile of nest extracts closely matches the ant’s CHC profile (see **Fig. S3** for the full chromatogram). **(B)** Experimental design for generating and testing CHC extracts. CHCs were extracted by immersing 300 ants in hexane for 10 minutes. Colonies of 100 ants and 50 pupae were introduced into arenas containing two patches, one treated with a cuticular extract and one with hexane (solvent control). **(C)** Percentage of observations in which colonies nested on the site treated with CHC extract, the hexane-treated site (solvent control), or neither. Colonies aggregated significantly more at sites treated with CHC extract than at sites treated with hexane or at untreated sites. N = 13 replicates. **, *adj. P* < 0.01. An example image of the experimental setup is shown on the right. **(D)** Location heatmaps showing the spatial distributions of 100 ants from the foraging phase, either young or old, over 1 minute in arenas in which CHC extract or hexane had been applied in the shape of the letter “P.”

Notably, when applied to the experimental arena in the form of an arbitrary shape, the cuticular extracts elicited the formation of corresponding unnatural spatial distributions of ants that lasted at least several minutes **(Fig. 3D, Video S1)**. The application of hexane in a similar shape failed to elicit this response **(Fig. 3D)**. This suggests that ants use CHCs transferred to the substrate to determine the location of the nest.

### Responsiveness to the arrestant effect of hydrocarbons correlates with the ants’ tendency to remain inside the nest

The tendency of *O. biroi* to remain inside the nest varies with both age and phase of the colony cycle. We therefore asked whether these behavioral differences are associated with differential arrestment responses elicited by CHCs **(Fig. 4A)**. By assaying individual ants, we asked whether cuticular extracts alone, independent of additional cues from nestmates, are sufficient to elicit behaviors associated with nest fidelity.

**Figure 4.**
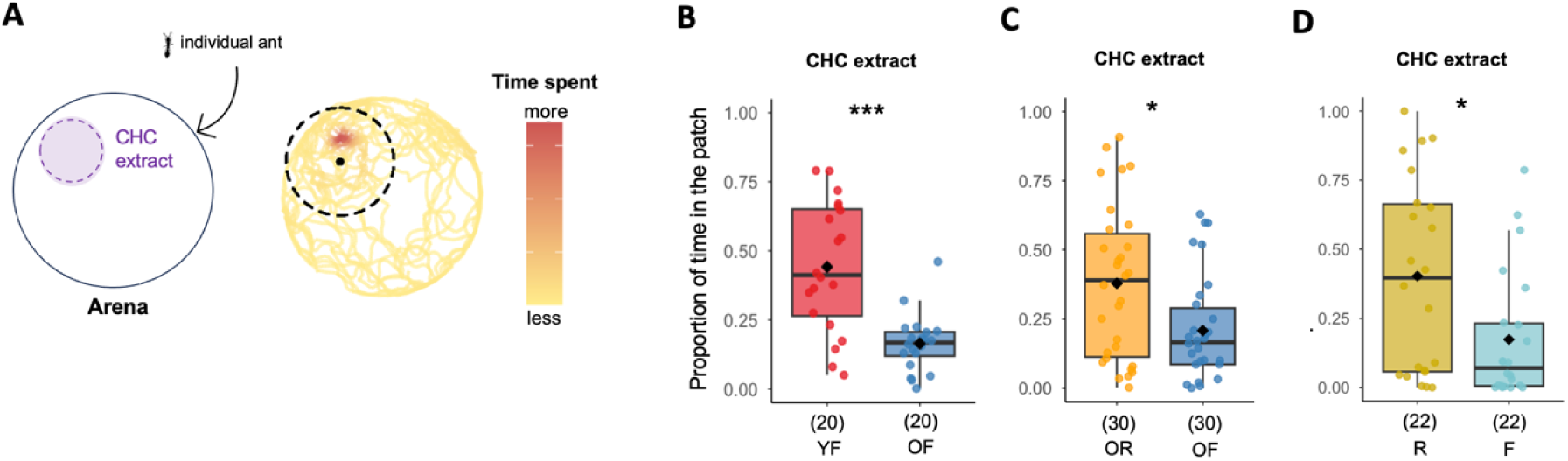
Responsiveness to the arrestant effect of hydrocarbons correlates with the ants’ tendency to remain inside the nest. **(A)** Left: Schematic of the assay to test individual arrestment in response to CHC extracts. Ants were introduced one-by-one into arenas containing a patch treated with CHC extract. Right: A representative trajectory of a young ant exploring an arena with a patch of CHC extract. The color scale indicates the amount of time spent in a given position during a 30-minute assay. The circle used to measure arrestment (proportion of time spent inside the circle) was slightly larger than the CHC extract application area to account for possible diffusion of CHCs (see Methods). **(B-D)** Proportion of time spent in the patches coated with CHC extract for different types of ants. The y-axis represents the proportion of the 30-minute test phase spent in the CHC-treated area. Each dot corresponds to an individual ant. Boxplots delineate the upper and lower quartiles, whiskers represent 1.5 times the interquartile range, diamonds indicate the mean, and horizontal bars the median. Sample sizes are reported on the x-axes. **(B)** Young ants (5-6 days old) from colonies in the foraging phase (YF, red) were more arrested by CHC extracts than old ants (4 months old) from colonies in the foraging phase (OF, blue). **(C)** Old ants (>1.7 months old) from colonies in the reproductive phase (OR, orange) were more arrested by CHC extracts than old yet slightly younger ants (> 1.2 months old) from colonies in the foraging phase (OF, blue). **(D)** 3–4-week-old ants in the reproductive phase (R, gold) were more arrested by CHC extracts than age-matched ants in the foraging phase (F, light blue). *, *P* < 0.05; ***, *P* < 0.001.

In each trial, a single ant was introduced into an arena containing a patch treated with either a CHC extract **(Fig. 4A)** or hexane **(Fig. S6A)**. After a 30-minute habituation phase, the ant location was tracked during a 30-minute test phase. To allow ants of different types (e.g., young vs. old or in reproductive phase vs. in foraging phase) to experience the same odor patch (either coated in CHC extract or hexane control), individuals were tested sequentially in the same arena (each arena was used for four individuals, see Methods) while alternating between types to minimize potential confounding effects from sensory cues left by other ants or gradual evaporation or diffusion of CHCs on the substrate.

Young ants were significantly more arrested by cuticular extracts than old ants, mirroring the stronger tendency of young ants to remain inside the nest (controlling for phase of the colony cycle, Wilcoxon rank-sum test*, P* < 0.001; **Fig. 4B**). Ants from colonies in the reproductive phase spent more time near cuticular extracts than ants from colonies in the foraging phase, again reflecting their greater tendency to stay in the nest (Wilcoxon rank-sum test, *P* < 0.05; ants from the reproductive phase were on average two weeks older than ants from the foraging phase in this comparison; **Fig. 4C**). To further separate age from colony-cycle effects, we repeated this assay using age-matched ants. We prepared replicate age-matched colonies with young ants (2-3 days old) and provided them with different brood types sourced from different stock colonies to induce transitions into different phases of the colony cycle^29^. After three weeks, we tested ants from these colonies in the same individual assay. This age-matched comparison confirmed that ants in the reproductive phase showed stronger arrestment by cuticular extracts than ants in the foraging phase (Wilcoxon rank-sum test, *P* < 0.05, **Fig. 4D**).

In contrast, the ants’ responses to the solvent control did not vary significantly across individuals of different ages or from different phases of the colony cycle, indicating that these behavioral differences do not generalize to different chemical compounds, or at least not to solvent residues (**Fig. S6B-C**). These findings suggest that behavioral responses to CHCs transferred to the substrate change with age and cycle phase, supporting a functional link between an ant’s responsiveness to the arrestant effect of CHCs and its tendency to aggregate and remain inside the nest in a colony setting.

### Hydrocarbons act as an aggregation pheromone

Pheromones are defined as chemical signals that elicit innate and stereotyped behavioral responses that are conserved across individuals and populations of a species. This distinguishes them from other chemical cues that are learned and elicit context-dependent behaviors^30^. For example, ants learn the CHC profile of their colony and use CHCs to distinguish nestmates from non-nestmates^31^. This is also the case in *O. biroi*, where CHC profiles differ between colonies of different clonal genotypes, providing salient cues for non-nestmate discrimination^32^ (**Fig. 5A, Fig. S7A, Table S1**). However, CHCs can act as pheromones in other contexts, for example, as fertility signals that regulate worker physiology and behavior^33^. To determine whether CHCs act as pheromones in the context of aggregation, we first asked whether the observed arrestment response reflects a general response to hydrocarbons, or whether it is specific to compounds found on the *O. biroi* cuticle. We then tested whether the arrestant effects of *O. biroi* CHCs generalize across different clonal genotypes.

**Figure 5.**
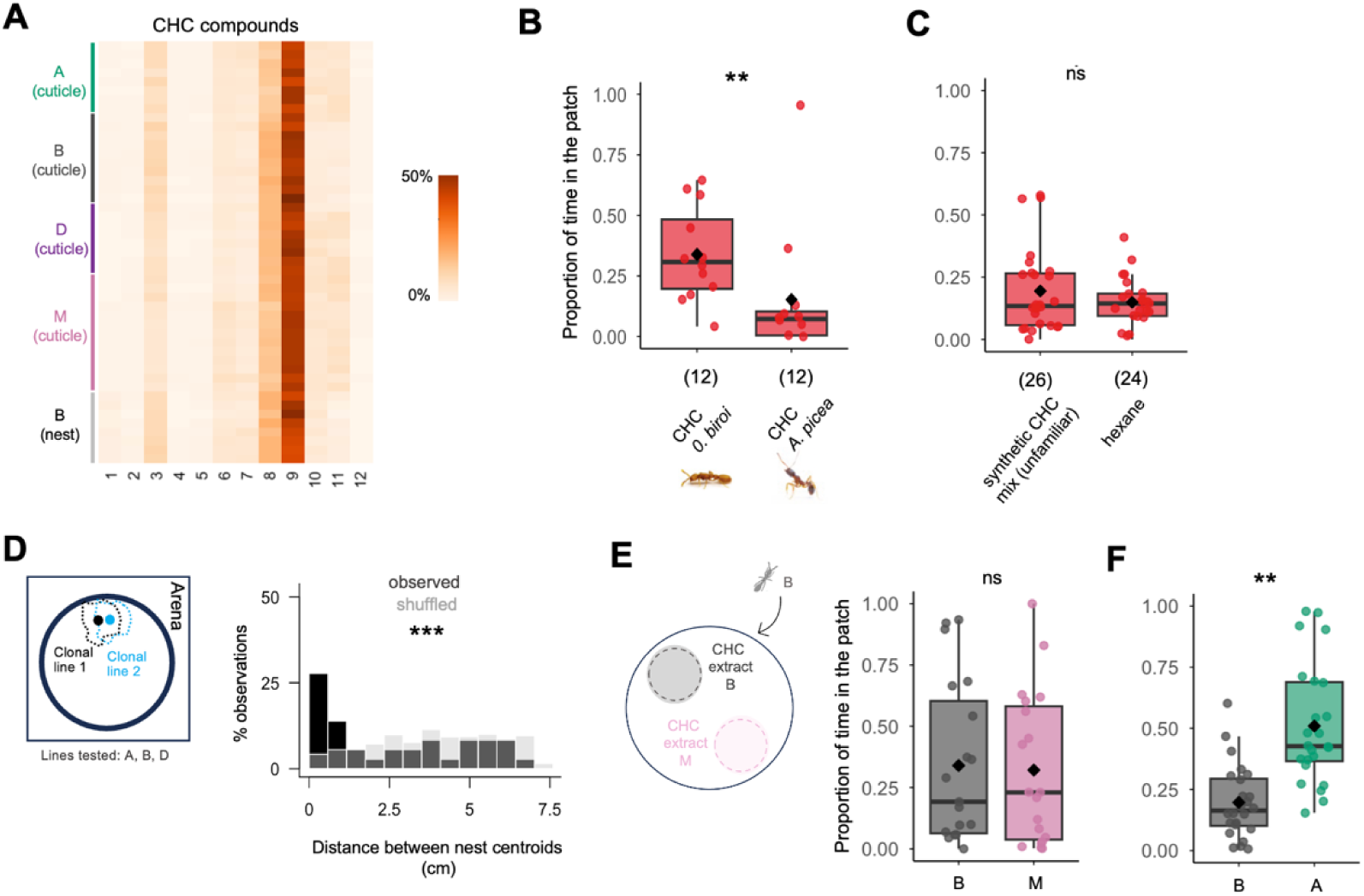
Hydrocarbons act as an aggregation pheromone. **(A)** Heatmap of hydrocarbon compounds from the cuticles of different *O. biroi* clonal lines and from nests of clonal line B. Each column represents an identified compound, and color intensity indicates the relative abundance on the cuticle or on the substrate (0-50% of total CHC mass). Several GC peaks contained co-eluting positional isomers of methyl-branched alkanes, which are listed together in the following key: 1 = C25, 2 = 13-;11-MeC25, 3 = 2-;3-MeC25, 4 = C26; 3,7-DiMeC25, 5 = 13-;12-MeC26, 6 = 2-;3-MeC26, 7 = C27, 8 = 13-;11-MeC27, 9 = 11,15-DiMeC27; 2-MeC27, 10 = 3-MeC27, 11 = 3,11-;3,9-;3,7-DiMeC27, 12 = 11,15-DiMeC28, 2-MeC28 (see also Table S1). (**B)** Young *O. biroi* workers (3-7 days old) in the foraging phase arrested more in response to *O. biroi* CHC extracts than in response to *Aphaenogaster picea* extracts. **(C)** A synthetic CHC blend composed of hydrocarbons absent from the *O. biroi* cuticle (13,23-DiMeC36 + 2-MeC30 + C30) did not elicit significant arrestment compared to hexane controls in young ants (3-7 days old) in the foraging phase. **(D)** Nest-site overlap assay for colonies of different genotypes (A, B, D) composed of 100 old ants in the reproductive phase and 50 pupae. Left: schematic illustrating the distance measurement between nest centroids of two colonies of different genotypes. Right: distribution of observed distances between nest centroids (black) compared with shuffled controls (light gray; nest positions permuted across independent experiments). Colonies nested significantly closer to sites previously occupied by colonies of a different genotype than expected by chance (36 replicates). **(E)** Left: Schematic illustrating the choice assay to test genotype specificity of CHC-mediated arrestment in 1-month-old clonal line B ants in the reproductive phase. Individual ants were introduced into arenas containing two patches, one treated with a cuticular extract from their own genotype (line B) and the other treated with an extract from a genetically distinct genotype (line M). Right: Arrestment was quantified as the proportion of the test phase spent within each treated area. Line B ants showed no significant difference in arrestment between extracts from line B and line M (20 replicates). **(F)** Same as (E), but comparing cuticular extracts from line B to extracts from line A. Line B ants spent significantly more time on line A than on line B extracts (24 replicates). Each dot represents one ant. Boxes indicate the interquartile range, whiskers represent 1.5 times the interquartile range, diamonds indicate means and horizontal lines indicate medians. ns, *P* > 0.05; **, *P* < 0.01.

Cuticular extracts from *Aphaenogaster picea*, a distantly related ant species, induced significantly less arrestment in young ants than *O. biroi* extracts (Wilcoxon rank-sum test, *P* < 0.01) (**Fig. 5B**).

To further test the specificity of the arrestment response, we tested a synthetic CHC blend composed of linear, methyl-branched, and dimethyl-branched alkanes that are structurally similar to *O. biroi* CHCs but absent from the *O. biroi* cuticle. Young ants showed no significant arrestment by this blend compared to hexane controls (Wilcoxon rank-sum test, P > 0.05; **Fig. 5C**). These results suggest that arrestment is not elicited by hydrocarbons in general, but that it depends on compounds specifically present in the *O. biroi* CHC profile instead.

Ants typically become better at discriminating nestmates from non-nestmates based on CHC profile differences as they get older^31^. We therefore conducted nest-site overlap assays using colonies with old ants and pupae, introducing them to arenas that had previously been occupied by *O. biroi* colonies of different genotypes (**Fig. 5D**, see Methods). Colonies preferentially established nests at previously occupied sites, even if the previous occupants were from a different clonal line (Wilcoxon rank-sum test, *P* < 0.001). This contrasts with the fact that ants from these different clonal lines behave antagonistically in direct encounters^32,34^.

To corroborate these findings, we introduced one-month-old ants from clonal line B colonies in the reproductive phase into arenas containing two CHC-treated patches: one treated with a cuticular extract from ants of their own genotype (clonal line B) and the other with an extract from ants of a different genotype (either line M or line A). Ants of this age and from these different clonal lines recognize each other as non-nestmates in direct interaction assays^32^ (**Fig. S7B**). We then quantified the proportion of time each ant spent in each patch. The ants did not prefer extracts from their own genotype over extracts from the other two genotypes (**Fig. 5E** (line M) and **Fig. 5F** (line A)). In contrast, clonal line B ants spent significantly more time on extracts from clonal line A ants (Wilcoxon signed-rank test, *P* < 0.01). Behavioral observations showed that this difference was due to normal arresting behavior, rather than aggressive behavior directed toward the patches with line A extract (**Video S2**). This is consistent with the fact that clonal line B ants are not more aggressive toward line A ants than toward ants from other clonal lines, including line M (**Fig. S7B**). Furthermore, ants of clonal line A tend to spend more time in the nest than ants of clonal line B^24^, raising the possibility that line A cuticular extracts elicit particularly strong arrestment responses when transferred to the substrate, and that this effect generalizes across different *O. biroi* clonal lines.

Together, these experiments show that chemical extracts from the *O. biroi* cuticle but not extracts from other ant species or synthetic mixtures of unfamiliar hydrocarbons promote arrestment. Extracts from one clonal line elicit arrestment in workers of different clonal lines. *O. biroi* cuticular extracts thus elicit stereotyped arrestment and aggregation responses across different *O. biroi* clonal lines, establishing them as aggregation pheromones when transferred to the substrate.

### A single hydrocarbon compound elicits aggregation

Having established that *O. biroi* cuticular extracts contain an aggregation pheromone, we wanted to determine which compound(s) mediate this effect. While CHCs have been implicated in a variety of social functions in ants^33,35,36,37^, their specific roles are rarely attributed to individual compounds or specific combinations of compounds. *O. biroi* might offer a useful system for dissecting this complexity. The species’ CHC profile is relatively simple, comprising a maximum of 32 compounds, but with only three major compounds (11,15-dimethylheptacosane, 13-methylheptacosane and 2-methylpentacosane) that account for approximately 70% of the total CHC mass in adults **(Fig. 5A**, **Fig. 6A)**.

**Figure 6.**
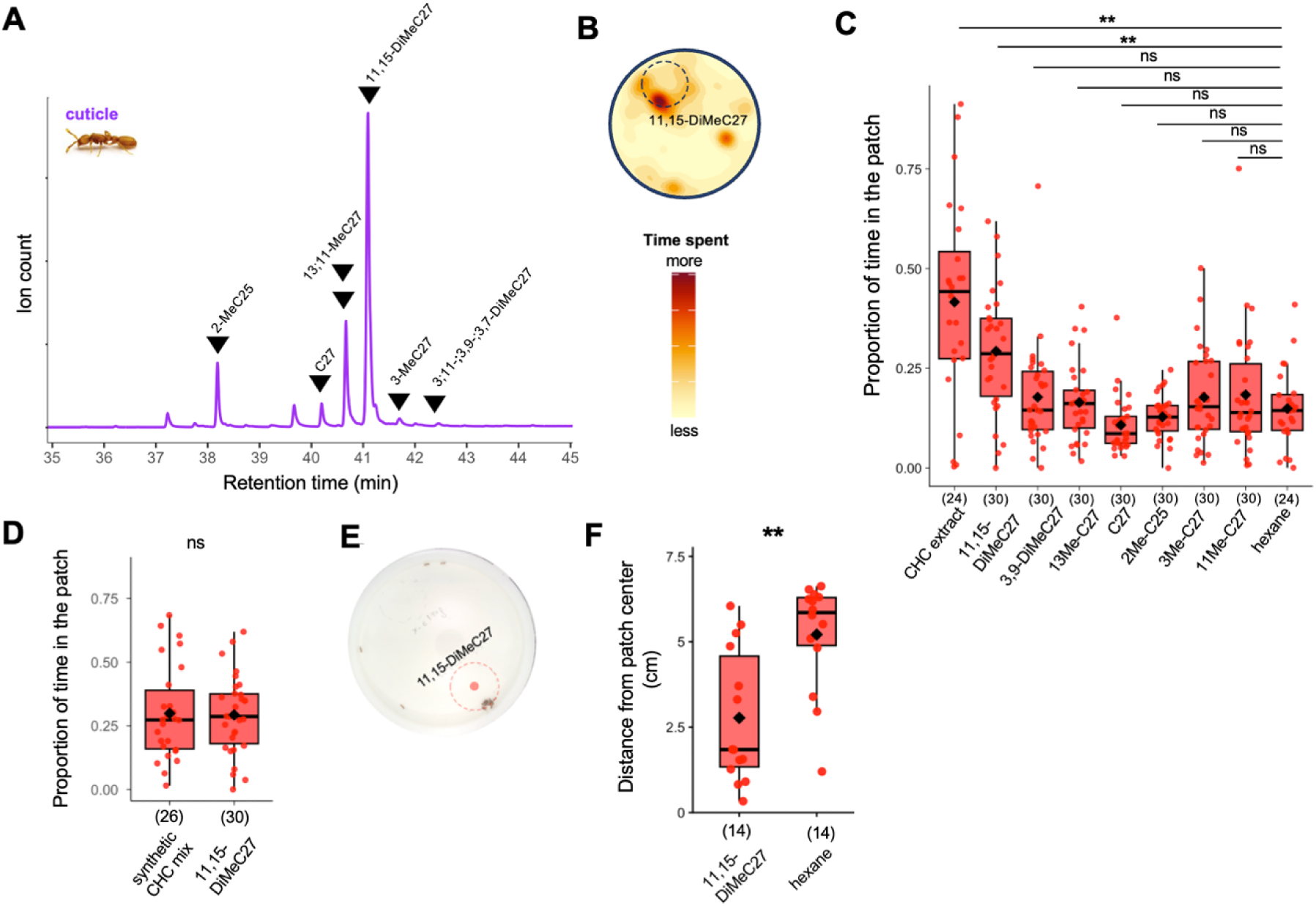
A single hydrocarbon compound elicits aggregation. **(A)** Representative chromatogram of cuticular extract from an *O. biroi* worker, showing the different CHC compounds. Black triangles indicate CHCs that were synthesized or commercially acquired and tested in behavioral assays. Among the possible configurations consistent with the fragmentation pattern for the peak corresponding to 3,11-;3,9-;3,7-DiMeC27, only 3,9-DiMeC27 was synthesized, whereas for 13;11-MeC27, both 13-and 11-MeC27 were synthesized (see Methods). **(B)** Example heatmap of an individual young ant (3-7 days old) in the foraging phase in an arena treated with 11,15-DiMeC27. **(C)** Behavioral responses of young ants (3-7 days old) in the foraging phase to synthetic CHC compounds. Individual ants were tested in arenas containing a patch treated with synthetic CHCs, CHC extract, or hexane solvent control. Arrestment was quantified as the proportion of time spent inside the treated area during a 30-minute assay. Among the tested compounds, 11,15-DiMeC27 elicited strong arrestment whereas other single CHCs produced no significant effects. Significance is shown relative to hexane (Wilcoxon rank-sum tests with Bonferroni–Holm correction; **, *adj. P* < 0.01; ns, *adj. P* > 0.05). **(D)** Young ants (3-7 days old) in the foraging phase were equally arrested by 11,15-DiMeC27 and by a synthetic mix of the three most abundant components of the *O. biroi* CHC profile in natural proportions (5:2:1 ratio of 11,15-DiMeC27, 13-MeC27, and 2-MeC25). ns, *P* > 0.05. Dots correspond to individual ants. **(E)** Example of a group assay showing ants aggregating near a patch treated with 11,15-DiMeC27. **(F)** Groups of young ants (20 ants per group; 3-7 days old) in the foraging phase aggregated on average closer to patches treated with 11,15-DiMeC27 than to patches treated with hexane. Each dot represents one group tested. N = 14 replicates for each treatment. **, *P* < 0.01. Boxes indicate the interquartile range, whiskers represent 1.5 times the interquartile range, diamonds indicate means and horizontal lines indicate medians. Sample sizes are reported in parentheses below the x-axis.

To identify specific *O. biroi* CHCs that mediate aggregation, we purchased or synthesized most of the most abundant compounds on the *O. biroi* cuticle (**Fig. 6A)** (see Appendix for synthesis methods). We then tested their behavioral effects in assays where we applied the compounds to featureless arenas lined with filter paper, introduced individual young ants, and monitored their behavior as described above (**Fig. 6B)**. We conducted these behavioral assays with young ants, because they showed the most robust aggregation responses (**Fig. 2C**). To assure that these assays were biologically relevant, the amounts of applied synthetic CHCs corresponded to extracts from ∼70-700 ants (**Fig. S8**, see Methods).

A single compound, 11,15-dimethylheptacosane (11,15-DiMeC27), was sufficient to induce arrestment in young ants (**Fig. 6C**). This compound, which is the most abundant hydrocarbon on the *O. biroi* cuticle and accounts for nearly 50% of the total CHC mass, elicited significantly greater arrestment than the solvent control (Kruskal-Wallis test P < 0.001; Wilcoxon rank-sum test with Bonferroni–Holm correction, *adj. P* < 0.01). None of the other six synthetic CHCs tested elicited significant arrestment when presented individually (**Fig. 6C**). We then tested a mix containing 11,15-DiMeC27 and the two other major compounds (13-MeC27 and 2-MeC25) at ratios that closely recapitulated the composition of the *O. biroi* CHC profile (see Methods). This synthetic blend did not elicit significantly greater arrestment than 11,15-DiMeC27 alone (Wilcoxon rank-sum test, *P* > 0.05; **Fig. 6D**). Finally, the effect of 11,15-DiMeC27 was also apparent in a group context: groups of young ants (n = 20) aggregated on average closer to patches treated with 11,15-DiMeC27 than to patches treated with hexane (Wilcoxon rank-sum test, *P* < 0.01) **(Figs. 6E-F)**. Taken together, these results show that 11,15-DiMeC27 serves as an *O. biroi* aggregation pheromone.

## Discussion

The evolution of eusociality in ants likely required numerous changes in sensory and neural systems that allowed stable group living. Identifying the cues that promote social cohesion among ant workers therefore provides a useful entry point into understanding how ants evolved their social lifestyle. Here, we show that cuticular hydrocarbons (CHCs), when transferred to the nest substrate, are sufficient to promote aggregation and to indicate the location of the nest. Although an ant’s decision to remain with nestmates is likely mediated by multiple factors, possibly including direct tactile interactions with other workers, our results show that substrate-borne chemical cues alone play an important role in eliciting aggregation behavior as part of a colony’s “social glue.”

The CHC-based pheromone described here differs from aggregation or recruitment pheromones that attract individuals from a distance (e.g.^10,38^). Instead, it acts at short range, promoting local arrestment and stabilizing aggregations once workers encounter a marked site. This effect is reminiscent of weakly volatile aggregation pheromones in termites and fruit flies^11,12,13^ (but see^14^). CHCs are known to accumulate in ant nests and other insect environments (e.g. ^39,40^), usually passively through repeated contact between the body surface and the substrate^41,42^. In ants, CHCs transferred to the substrate can serve to mark territories^43^ and nest entrances^44,45^, and to spatially organize the nest to regulate division of labor^36^. Similarly, our findings show that, in *O. biroi*, CHCs transferred to the nest surface are not merely residual byproducts of occupancy but have important functions as aggregation-promoting signals.

In *O. biroi*, as in other ants, differences in CHC profiles between colonies provide salient cues for non-nestmate discrimination^32,34^. In this context, CHC cues are learned and interpreted relative to prior social experience^31,32,46^. In contrast, the aggregation-promoting activity we identify here generalizes across different colonies and different clonal genotypes, and the behavioral response is likely innate. These properties imply that CHCs, when transferred to the substrate, function as a pheromone during nest aggregation. CHCs therefore function in at least two different contexts in *O. biroi*: as colony-specific cues relevant to non-nestmate discrimination^32^ and as pheromones that promote group cohesion (this study). Our results further suggest that CHCs on the substrate are sufficient to drive aggregation behavior but insufficient to elicit non-nestmate discrimination in the absence of additional sensory cues. This in turn implies that the behavioral response to CHCs depends on the sensory context.

The most abundant CHC on the *O. biroi* cuticle, 11,15-dimethylheptacosane (11,15-DiMeC27), by itself was sufficient to elicit arrestment in young workers, implying that the core signal that promotes aggregation is remarkably simple. Cuticular profiles in ants are typically complex and multifunctional, and specific behavioral functions are rarely assigned to individual CHCs^26,27,33,47^, making this finding notable.

At the same time, the synthetic compounds tested here did not fully reproduce the arrestant effect of complete cuticular extracts (**Fig. 6C**), suggesting that additional factors might contribute to aggregation. For example, minor CHC components not included in our synthetic blends may enhance arrestment, or cuticular extracts may contain small amounts of non-long-chain hydrocarbon compounds that contribute to the behavioral effect. Alternatively, the synthetically produced compounds might differ from the native compounds in ways that affect behavior. For example, the synthetic dimethyl-branched hydrocarbons used here were racemic mixtures (see SI Appendix), whereas natural CHCs likely occur in only one stereochemical form^48^. This difference could affect receptor binding and perceptual valence.

Finally, the ants’ responsiveness to substrate-borne hydrocarbons varied with worker age and phase of the colony cycle in a way that paralleled their natural tendency to remain inside the nest. Young workers, which typically take on the role of nurses, arrested more in response to aggregation pheromones than older workers, which typically forage. Workers from colonies in the reproductive phase, which do not forage, likewise showed stronger responses to aggregation pheromones than workers from colonies in the foraging phase. This suggests that the neural and behavioral responses to aggregation pheromones are plastic, which in turn contributes to the spatial organization of behavior and the division of labor in ant colonies. This is consistent with previous studies reporting a role of substrate-borne CHCs in the spatial segregation of different worker castes in *Pogonomyrmex* harvester ants^35^ and *Camponotus* carpenter ants^36^.

The roles of CHCs as substrate-borne cues in eusocial insects might be ancient and even predate the origin of eusociality. Ant eusociality is thought to have evolved from subsocial ancestors with progressive provisioning, a form of parental care in which individuals repeatedly return to a stable nest to feed their larvae^49,50^. Extant progressive provisioners transfer CHCs to the substrate around the nest entrance, and they might use these compounds as olfactory signposts when returning home^41^. This ancestral arrestment response to substrate-borne CHCs might have predisposed offspring to remain at their mother’s nest, a major step in the evolution of eusociality^49,50^. Finally, if the plasticity in behavioral responses to substrate-borne CHCs we observe in *O. biroi* also dates back to the origin of ant eusociality, it could have immediately contributed to emergent division of labor in incipient eusocial groups along the axes of age and reproductive status. Additional studies across a broader taxonomic range of eusocial and subsocial taxa will be required to reconstruct these evolutionary trajectories more fully.

## Materials and Methods

Materials and Methods are provided in the Supplementary Information.

## Data and code availability

Relative-abundance datasets for CHCs from different developmental stages, adult colony phases, clonal genotypes, and nest extracts are provided in Datasets S1-S3. Behavioral data and analysis code are available on Github (https://github.com/MatteoRossiRU/Obiroi_aggregation).

## Author Contributions

M. R., T.P.M.B., E.T.F. and D.J.C.K. designed research with input from J.G.M., S.V-R. and T.S.; M.R., T.P.M.B. and E.T.F. performed research; J.G.M. and D.O.M. contributed new reagents/analytic tools; M.R., T.P.M.B. and E.T.F. analyzed data; M.R. and D.J.C.K. wrote the manuscript with input from all authors; M.R. prepared figures with input from D.J.C.K. D.J.C.K. supervised the project.

## Artificial Intelligence (AI) Disclosure

AI software was used for manuscript editing (ChatGPT-6; Claude Sonnet 4.6 – 5.5) and analysis code development (ChatGPT-6). The authors have thoroughly checked all edits to both the manuscript and analysis code, and take full responsibility for the final products.

## Acknowledgments

We thank Tomas Kay, Libby Tseng, Yannick Günzel and Yukina Chiba for comments on an earlier version of this manuscript. This work was supported by a European Molecular Biology Organization Fellowship (ALTF 955-2022) and a Human Frontier Science Program Fellowship (LT0023/2023-L) to M.R. T.P.M.B. was supported by a European Molecular Biology Organization Fellowship (ALTF 131-2022), a Leon Levy Scholarship in Neuroscience, and a fellowship from the Price Family Center for the Social Brain. Additional support was provided by the Howard Hughes Medical Institute, where D.J.C.K. is an investigator. J.G.M. was supported by Hatch Act project CAR*ENT-5181-H. T.S. was supported by the University of Würzburg. E.T.F. was supported by the Emmy Noether Program of the German Research Foundation (DFG) (grant number: 511474012). This is Clonal Raider Ant Project paper #46.

## Supplementary Information

### Materials and Methods

#### Ant husbandry

Clonal raider ant (*Ooceraea biroi*) colonies were maintained in environmental chambers at 25°C and 60% relative humidity in plastic containers with a moist plaster of Paris floor. During the foraging phase, colonies were fed frozen fire ant brood on a regular schedule (every Monday, Wednesday and Friday). For most experiments, ants were derived from laboratory stock colonies of clonal line B (originally collected on Saint Croix, U.S. Virgin Islands in 2009). Some experiments also used ants from clonal line A (originally collected on Okinawa, Japan in 2008), as well as clonal lines D and M (originally collected in Bangladesh in 2014). One experiment used *Aphaenogaster picea* workers (collected at the Rockefeller University Field Center in Millbrook, NY, in 2024, and maintained in a terrarium with a substrate of soil, twigs, and bark fragments from the collection site).

#### Experimental arenas

Behavioral assays were performed in laser-cut circular acrylic arenas (76 mm internal diameter). The arenas were covered with a lid and fitted with a fresh plaster of Paris base. The plaster base was lined with a sheet of 0.18 mm thick Tyvek® (DuPont) and then covered with white filter paper (Fisher Scientific, Cat. No. 09-801E, 09-803-6D). The plaster base was hydrated with distilled water before each experiment. To prevent carryover of chemical cues between trials, the plaster base, filter paper and Tyvek were replaced after each behavioral assay. Acrylic parts were soaked overnight in detergent, rinsed thoroughly with water, air-dried, and wiped with 96% ethanol before reuse.

#### Nest-site overlap assays

Experimental colonies consisted of 100 old workers in the reproductive phase and 50 pupae. Ants and pupae were sourced from large stock colonies, where they were picked up with soft forceps and transferred into Petri dishes. They were then scattered across the experimental arenas by gently tapping the Petri dish. A first colony was introduced into a fresh arena and allowed to settle for 16–20 hours under continuous video recording. The colony was then removed by transferring the ants and pupae into a Petri dish with a brush, and a second, naïve colony of identical composition was introduced either 5 minutes or 24 hours later. To exclude the possible influence of cues external to the arena on nest site choice, the arena was rotated by 90° between removal of the first colony and introduction of the second colony. The second colony was recorded for 1 hour following introduction. A total of 12 colonies of clonal line B were used as biological replicates. Each colony was tested in three independent technical replicates under each of the two interval conditions (5 minutes and 24 hours). Across these trials, each colony was introduced first and second six times each. The two interval conditions were tested in alternating order (**Fig. 1D**).

For assays with a 24-hour interval before introducing the second colony (**Fig. 1D**), the Tyvek sheet and filter paper lining the arena floor were removed from the arena (after the first colony was taken out). The Tyvek and filter paper were then stored in a sealed container at room temperature for 24 hours to permit evaporation of chemical cues. Before introducing the second colony, the plaster base was rehydrated, and the original Tyvek sheet and filter paper were refitted to the arena.

To test whether nest-site overlap depended on colony genotype, we performed the same assay using colonies from clonal lines A, B, and D that were synchronized in their colony cycle phase by setting up colonies simultaneously without brood (**Fig. 5D**). Colonies consisted of 100 old workers in the reproductive phase and 50 pupae. Colonies were allowed to establish a nest for 16–20 hours before removal, after which a second colony of a different clonal genotype was introduced following the same procedure described above, with a 5-minute interval between colonies. All pairwise combinations of clonal lines A, B, and D were tested. Four colonies were used as biological replicates for each clonal line. For each genotype combination, each colony was introduced first and second three times each.

To determine whether the presence of pupae influenced nest-site overlap between colonies (**Fig. S1**), we compared colonies of 100 old workers in the reproductive phase containing 50 pupae with matched colonies without pupae (sourced from the same clonal line B stock colonies) using the same assay. Because colonies without brood gradually dispersed over time in pilot experiments, colonies with and without pupae were both allowed to settle for only 3 hours in the arena before removal of the first colony and introduction of the second. This shorter interval allowed us to quantify nest aggregation before substantial disaggregation occurred in groups without pupae, while maintaining identical conditions across treatments. For the condition with pupae and without pupae, 36 and 35 colonies were used as biological replicates, respectively, and each colony was only used once. Each colony was introduced first and second once each.

#### Colony cohesion assays

To quantify colony cohesion (**Fig. 2B-C**), we assembled three types of experimental groups consisting of 100 workers each: young workers (3–7 days old) from clonal line B stock colonies in the foraging phase, old workers (>1.2 months old) from clonal line B stock colonies in the foraging phase (collected from the same stock colonies as the young workers), and old workers (>1.7 months old) from stock colonies in the reproductive phase. Groups were introduced into experimental arenas and allowed to settle for 3 hours under the same conditions as described above, before colony cohesion was quantified (see below). For each of the three conditions, 20 groups were used as biological replicates.

#### Automated quantification of nest location and colony cohesion

Arenas were imaged from above at 5 fps or 10 fps and 2,592 × 1,944-pixel resolution using webcams (Logitech C910). Videos were analyzed with a custom MATLAB pipeline. To estimate the nest location, images were converted to grayscale and normalized to a 0–1 intensity scale (0 = black, 1 = white). Pixels with grayscale intensity < 0.5 were classified as foreground, corresponding to dark pixels associated with ant bodies. The square arena image was divided into a 6 × 6 grid, and a square sliding window corresponding to one grid cell was moved across the arena in increments of 20% of the window width. The number of foreground pixels within each window was quantified, and nest position was defined as the centroid of the window containing the greatest number of foreground pixels (corresponding to ant bodies).

For nest-site overlap assays (**Fig. 1D, Fig. S1, Fig. 5D**), we analyzed the final 30 minutes before removal of the first colony and the final 30 minutes of the 1-hour recording of the second colony, sampling one frame every 30 seconds. For each frame, the group centroid was estimated from the foreground pixels as described above, and the centroid coordinates were averaged across the sampled frames to obtain a mean nest centroid. The Euclidean distance between the mean nest centroids of the first and second colony was calculated in pixels and converted to centimeters. To determine whether the observed distance between nest locations exceeded random expectations, null distributions were generated by estimating the distance between the nest centroids of the first and second colonies from independent arenas/experiments within the same treatment across all possible permutations. For each distribution, we calculated the number of observations falling within a given distance bin divided by the total number of observations in the corresponding distribution, multiplied by 100 (% observations).

For colony-cohesion assays (**Fig. 2B-C)**, a cohesion index was calculated as the proportion of foreground/dark pixels in the arena falling within the nest window. The index was calculated from the final 10 frames of the 3-hour recording, sampled every 30 seconds (corresponding to the final 5 minutes of the assay), and averaged across frames.

#### Extraction of chemicals from substrates

To collect chemical cues transferred to the nest substrate (**Fig. S2A-B**), three groups of 300 adult ants in the reproductive phase and 150 pupae were introduced into arenas prepared as described above and allowed to establish nests for 24 hours. After this, the ants and brood were removed, and two regions of the filter paper floor were sampled from each arena: a nest region occupied by the colony and a similarly sized region from an area of the arena that had remained unoccupied. For each extraction replicate, nest and non-nest (control) patches from the three independent arenas were pooled.

Filter-paper samples were immersed for 2 hours in 600 µL of one of five solvents spanning a range of polarities: hexane, dichloromethane, acetonitrile, ethanol, and methanol.

Dichloromethane should dissolve the hydrocarbon fraction but also more polar compounds. Why we did not observe robust behavioral responses to dichloromethane extracts is unclear. Acetonitrile, ethanol and methanol should not dissolve the hydrocarbon fraction or only dissolve minute amounts.

#### Behavioral assays with substrate extracts

After extraction, the solvent extract was recovered, and “nest” and “non-nest” extracts were applied to a filter paper for behavioral testing in the arenas. Solvents were allowed to evaporate under a fume hood before filter papers were fitted to the arenas. Evaporation times were 15 minutes for hexane and dichloromethane, 20 minutes for acetonitrile, and 30 minutes for ethanol and methanol. After fitting the filter paper to an experimental arena, we introduced a group of 100 workers (mixed age) from clonal line B in the reproductive phase with 50 pupae and recorded their behavior for 1 hour (**Fig. S2A-B**). Each nest extract and non-nest extract was tested only once. The nesting location of each colony was visually assessed from the last frame of the recording. Each independent behavioral assay was scored according to whether the colony nested on the nest extract, the control (non-nest) extract, or elsewhere. We calculated the percentage of observations for each outcome by dividing the number of assays assigned to that outcome by the total number of assays for the corresponding treatment and multiplying by 100. Statistical comparisons between nest and non-nest extracts for each solvent were considered exploratory, as the aim of this experiment was to identify potentially suitable solvents for subsequent chemical analyses. Therefore, P values were not adjusted for multiple comparisons. Behavioral data obtained with extracts from the two high-polarity solvents, ethanol and methanol, were pooled for analysis (15 and 16 replicates, respectively).

#### Gas Chromatography-Mass Spectrometry

Ants, pupae, and larvae were frozen at −20 °C prior to chemical extraction. Cuticular extracts were obtained from single adult ants in the foraging phase (3 months old) and reproductive phase (3.5 months old) from the same stock colony (clonal line B) (**Fig. S5B**). Cuticular extracts were also obtained from single adult ants of clonal line M in the foraging phase (1.5 months old) (**Fig. S7A**). Data for clonal lines A, B and D used for CHC profile comparisons were retrieved from^1^. Additional cuticular extracts were obtained from pooled samples of 100 larvae (3^rd^ and 4^th^ instars) and 100 pupae (1–2 days after pupation), derived from the same clonal line B stock colony (**Fig. S5A**). Cuticular hydrocarbons were extracted by immersing ants or brood in hexane for 10 minutes.

Nest and non-nest area extracts for GC-MS analysis were obtained by extracting a pool of filter paper pieces from 3 different arenas in hexane for two hours (**Fig. 3A**, **Fig. 5A**). The pieces of filter paper came from arenas in which 300 adult ants (mixed age, clonal line B) in the reproductive phase and 150 pupae had been allowed to nest for 2.5 days. We obtained additional extracts from nest and control areas using the same procedure from 3 different arenas that had housed either 300 young ants (4-8 days old, clonal line B) in the foraging phase or 300 old (>1.7 months old, clonal line B) ants in the reproductive phase, without any brood (**Fig. S4)**.

Extracts were evaporated to approximately 15 µL under a nitrogen flow, and 1 µL was injected into a 6890 GC unit coupled to a 5975 mass selective detector (Agilent Technologies) via a slit/splitless injector in the splitless mode at 300°C. The GC was equipped with a DB-5 capillary column (0.25 mm ID × 30 m; film thickness 0.25 μm). Helium was used as the carrier gas at a flow rate of 1 mL/minute. The temperature program started at 60°C, increased to 300°C at 5°C/minute, and was held at 300°C for 10 minutes. Mass spectra were recorded in EI mode at 70 eV with a source temperature of 230°C.

GC-MS data were analyzed using MassHunter Qualitative Analysis Navigator v.B.08.00 (Agilent Technologies). CHCs were identified using diagnostic ions and retention indices^2^. Retention indices were calculated by injecting commercially purchased n-alkanes. The relative abundance of each CHC compound was calculated as the integrated peak area divided by the sum of all CHC peak areas. Absolute CHC abundance was quantified by adding 10 µL of C18 (10 ng/μL; 100 ng total) to 15 samples of 2-week-old individual ants of clonal line B (C18 is not naturally present on the *O. biroi* cuticle) (**Fig. S8)**. Across adult cuticular extracts, nest extracts and brood extracts, we identified 32 hydrocarbon compounds, reported in Datasets S1 to S3.

In some cases, peaks contained partially co-eluting compounds. One peak was dominated in abundance by 11,15-dimethylheptacosane (11,15-DiMeC27) but also contained smaller amounts of 2-methylheptacosane (2-MeC27). To estimate the relative abundance of these two compounds when unresolved, we used diagnostic ions at *m/z* 351 and 239, which tracked their relative abundance in samples where the compounds separated clearly. A second peak was consistent with a mixture of 13-methylheptacosane (13-MeC27) and 11-methylheptacosane (11-MeC27).

Based on diagnostic ions at *m/z* 196 and 224 (characteristic of 13-MeC27) and at *m/z* 168 and 252 (characteristic of 11-MeC27), we found that these compounds were present at a 3:1 ratio across adult samples. Based on this consistent ratio, 13-MeC27 was treated as the predominant compound of the two isomers when making a synthetic CHC mix intended to approximate the natural *O. biroi* profile (see below). A third peak yielded a fragmentation pattern consistent with multiple 3,X-dimethylheptacosane configurations (shared diagnostic ions at *m/z* 56, 378, 408): 3,11-DiMeC27 (diagnostic ions at *m/z* 182 and 252), 3,9-DiMeC27 (*m/z* 154 and 280), and 3,7-DiMeC27 (*m/z* 126 and 308). Among these, the 3,9-configuration was the most abundant and was therefore selected for chemical synthesis and for individual-ant arrestment assays (see below). Finally, although 2- and 3-methylalkanes of the same chain length are expected to elute as separate peaks given their different retention indices, peak tailing caused them to partially overlap in some samples. Because these pairs could not be reliably integrated separately in all samples, we summed their peak areas and treated each pair as a single variable across all samples to ensure consistent quantification. These pairs are indicated with asterisks in Table S1 and Datasets S1-S3.

#### Group assays with cuticular extracts

Cuticular hydrocarbons were extracted by immersing 300 *O. biroi* workers in 400 µL of hexane for 10 minutes in a glass vial. Solvent-only controls consisting of 400 µL hexane were prepared in parallel. CHC extracts and/or hexane were applied to filter paper in one of four quadrants within the behavioral arena and allowed to evaporate for 15 minutes under a fume hood before testing. The location of the patch was alternated between the four quadrants of the arena to control for the possible effect of cues external to the arena on ant behavior.

In a first experiment (**Fig. 3B**), we applied hexane and CHC extracts to different patches in the same arena, introduced 100 ants (mixed age, clonal line B) in the reproductive phase with 50 pupae, and visually inspected where the colony had nested after 1 hour. Each independent behavioral assay was scored according to whether the colony nested on the CHC extract, the hexane patch, or elsewhere. We calculated the percentage of observations for each outcome by dividing the number of assays assigned to that outcome by the total number of assays and multiplying by 100 (**Fig. 3C**).

In a second experiment (**Fig. 3D)**, we prepared CHC extract by immersing 600 ants in 800 µL hexane and applied it in the form of the letter “P” to filter paper that was later fitted to a behavioral arena. We first introduced a group of 100 young ants (3-4 days old, clonal line B) in the foraging phase, recorded their behavior for 1 hour, and then introduced a group of 100 old ants (>1.2 months old, clonal line B) in the foraging phase for 1 hour into the same arena. In parallel, we applied 800 µL hexane in the shape of the letter “P” to another arena, where we introduced the same group of young ants (after having tested them with the CHC extract). For this experiment, ants were segmented and tracked using the TrackR software^3^.

#### Individual assays with cuticular extracts

To compare ants of different ages and from different cycle phases with respect to their arrestment response to CHCs or hexane (**Fig. 4, Fig. S6**), individual ants (clonal line B) were tested sequentially in the same arena. Four ants were assayed sequentially per arena, which contained a patch treated either with CHCs or hexane, in alternating order (for example, young–old–young–old or old–young–old–young). In age-comparison assays, young workers (5–6 days old) and old workers (4 months old) were obtained from the same stock colony.

To compare ants from different phases of the colony cycle in a first experiment (**Fig. 4C)**, we used workers (> 1.7 months old) sourced from colonies in the reproductive phase and workers (>1.2 months old) sourced from colonies in the foraging phase. Four individuals were tested in alternating order in the same arena as described above.

In a second experiment (**Fig. 4D)**, we prepared 4 colonies with 60 young ants each (2-3 days old, from the foraging phase), sourced from the same stock colony. We provided two of these colonies with 30 prepupae each and two with 30 larvae (2^nd^ instar) each. This treatment shifts initially identical ants into different phases of the colony cycle^4^. Brood items were sourced from different stock colonies than the one from which the young ants came. Individual ants from these age-matched colonies in different phases of the cycle were tested 3 weeks later (when they were 3-4 weeks old), again in alternating order in the same arena (4 individuals per arena, 2 individuals per ant type), which only contained a CHC-treated patch in this case.

In all these experiments, the individual ants were first allowed to explore the arena for 30 minutes (habituation phase), and we then recorded them for 30 minutes (test phase). The ants’ spatial coordinates throughout the test phase were recorded using anTraX^5^. The patch of interest was defined as a circular zone (same size across all experiments; radius = 2 cm, accounting for possible diffusion of the chemical stimulus on the filter paper) centered on the site of stimulus application. The proportion of time spent within this zone during the test phase was used as a measure of arrestment.

#### Individual assays with cuticular extracts from different species

To compare the arrestment response of young *O. biroi* ants to *O. biroi* and *Aphaenogaster picea* CHC extracts (**Fig. 5B**), we prepared CHC extracts from 300 *O. biroi* workers as described above, as well as from 50 *A. picea* workers (scaled to account for body size differences), by immersing ants for 10 minutes in 400 µL hexane. Young *O. biroi* ants (3-7 days old, foraging phase) were sourced from the same stock colony of clonal line B and were tested in different arenas, containing either a patch treated with cuticular extracts from *O. biroi* or *A. picea*. In each arena, we tested two young ants sequentially.

#### Individual assays with cuticular extracts from different *O. biroi* clonal lines

To compare the arrestment response of *O. biroi* to CHC extracts from different *O. biroi* clonal lines (**Fig. 5E-F**), we tested individual ants (1 month old, reproductive phase) from clonal line B in arenas that contained two different patches treated with either: i) CHC extracts from 300 individuals of clonal line B and clonal line M ii) CHC extracts from 300 individuals of clonal line B and clonal line A. In each arena, we tested four ants sequentially. Arrestment was quantified as described above, with the proportion of time spent within the circular zone surrounding each of the two stimulus patches calculated separately during the 30-minute test phase.

#### Individual assays with synthetic hydrocarbon compounds

We tested individual young workers (3–7 days old, clonal line B) in the foraging phase to quantify their arrestment response to individual synthetic CHC compounds, synthetic CHC mixtures, cuticular extracts, and hexane as a solvent control (**Fig. 5C**, **Fig. 6C-D**). For each assay, a single chemical stimulus was applied to one randomly selected quadrant of the arena. Two ants were tested sequentially in each arena, with each ant tested only once and exposed to only one stimulus at one dose. These assays were conducted using ants sourced from the same stock colonies, with the stimulus tested in each arena randomized across assays. Arrestment was quantified as the proportion of time each ant spent within the circular zone surrounding the stimulus application site during the 30-minute test phase, following the procedure described above for individual assays.

We tested each of the seven synthetic CHC compounds (**Fig. 6A**) individually at three doses: 0.05, 0.15, and 0.5 mg, with 10 ants tested per compound at each dose. These amounts correspond approximately to 70, 215, and 715 ant equivalents, respectively, based on an average yield of 700 ng of CHCs per ant cuticular extract (**Fig. S8**). For each compound, arrestment responses were compared across doses using Kruskal-Wallis tests. No significant dose effect was detected for any compound (all *P* > 0.05). We therefore pooled observations across the three doses for each compound for subsequent analyses, yielding 30 ants tested per individual synthetic CHC compound (**Fig. 6C**).

We also tested two synthetic CHC mixtures (**Fig. 5C**, **Fig. 6D**), each at a single dose of 0.24 mg, corresponding to approximately 340 ant equivalents based on the average CHC yield per ant (**Fig. S8**). The first mixture contained the three major components of the *O. biroi* cuticular profile, consisting of 0.15 mg 11,15-dimethylheptacosane (11,15-DiMeC27), 0.06 mg 13-methylheptacosane (13-MeC27), and 0.03 mg 2-methylpentacosane (2-MeC25). The three compounds were mixed in a 5:2:1 mass ratio, similar to the ratio in which they occur on the *O. biroi* cuticle. The second mixture contained hydrocarbons absent from the *O. biroi* cuticular profile: 0.15 mg 13,23-dimethylhexatriacontane (13,23-DiMeC36), 0.06 mg 2-methyltriacontane (2-MeC30), and 0.03 mg triacontane (C30) (0.24 mg total, 5:2:1 mass ratio). We tested a total of 26 ants per CHC mixture.

Finally, we tested 24 ants with *O. biroi* cuticular extracts (from 300 ants, as described above) and 24 ants with hexane as positive and negative controls, respectively (**Fig. 5C**, **Fig. 6C**). The ant-equivalent estimates do not account for the stereoisomeric composition of the synthetic compounds. In particular, the racemic mixtures of the dimethylalkanes should contain approximately 25% of the naturally occurring stereoisomer, and racemic mixtures of monomethylalkanes should contain approximately 50% of the naturally occurring compound.

However, this limitation is unlikely to substantially affect the interpretation of our results given that compounds were tested across a range of doses and no significant dose-dependent differences in arrestment responses were detected. Moreover, the biological activity of the alternative stereoisomers remains largely unknown. Although ants likely produce a single stereoisomer, this does not necessarily imply that their receptors are stereoselective. Other stereoisomers could potentially interact with the same receptors and elicit similar behavioral responses. Indeed, a study in the German cockroach reported that the naturally occurring stereoisomer of a dimethyl-branched sex pheromone elicited weaker courtship responses than the other three stereoisomers at physiologically relevant concentrations^6^.

#### Group assays with 11,15-dimethylheptacosane

Groups of 20 young workers (3-7 days old, clonal line B) in the foraging phase were released into an arena in which a patch of either 11,15-dimethylheptacosane (11,15-DiMeC27; 0.1 mg) or hexane (solvent control) had been applied to a randomized quadrant (**Fig. 6E-F**). Each group was only tested once and in a single condition.

Each assay lasted 30 minutes. For analysis, we examined the final 5 minutes of each assay, sampling one frame every 30 seconds. In each frame, the group centroid was estimated from the foreground/dark pixels corresponding to ants as described above, and the centroid coordinates were averaged across the sampled frames to obtain a mean group centroid. We then calculated the Euclidean distance between this mean centroid and the center of the stimulus application site.

### Supplementary Figures

**Figure S1.**
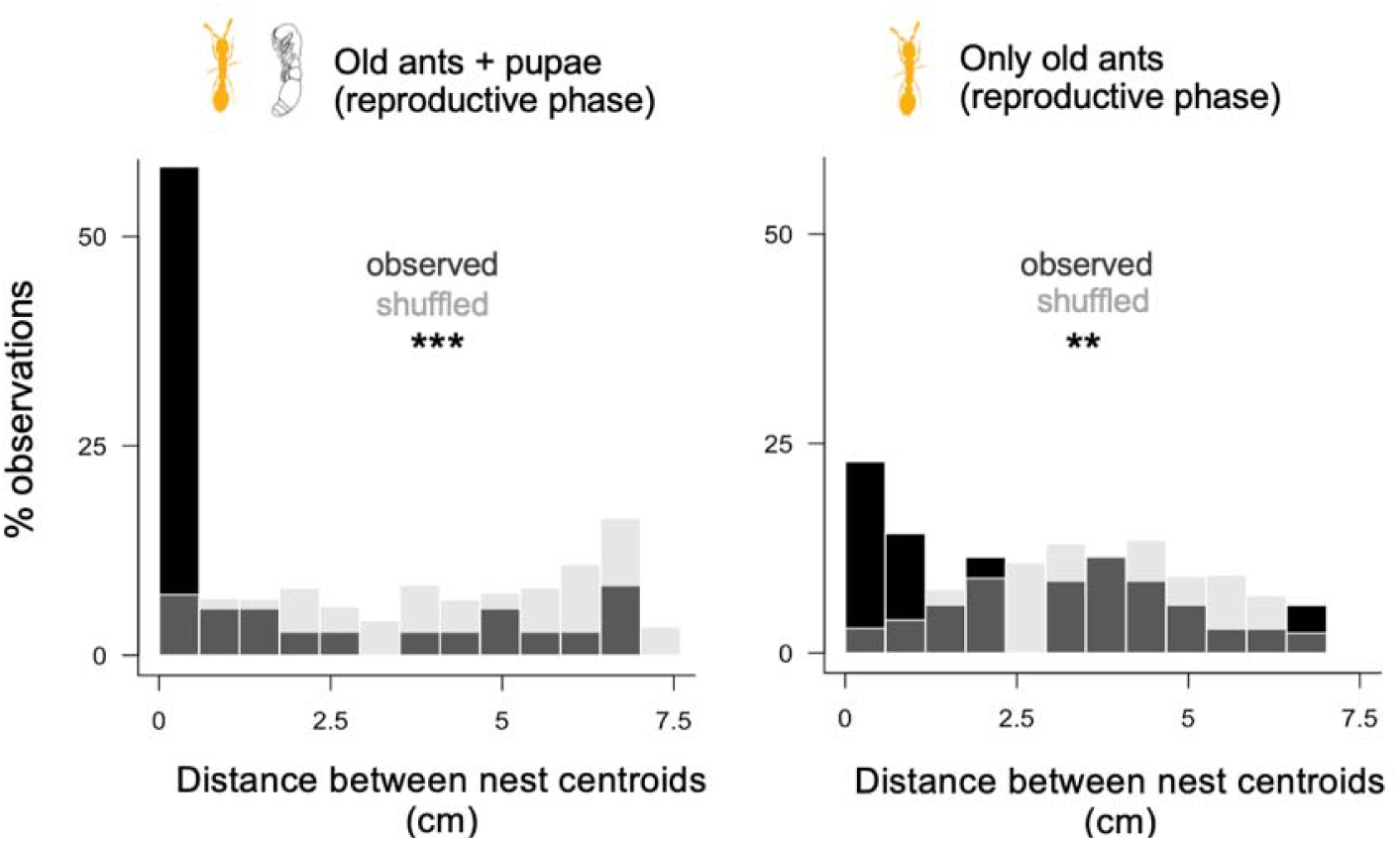
Adults are sufficient to mark the nest with an arrestant cue. Distribution of distances between successive nests for colonies in the reproductive phase with pupae (100 old adults with 50 pupae) or without brood (100 old adults only). The time elapsed between removing colony 1 and adding colony 2 was 5 minutes for each condition. Colonies with brood consistently reestablished nests near previous sites. Adults alone showed a weaker but still significant tendency to settle at previously occupied sites. **, *P* < 0.01; ***, *P* < 0.001. N = 36 and 35 replicates for the conditions with pupae and without brood, respectively.

**Figure S2.**
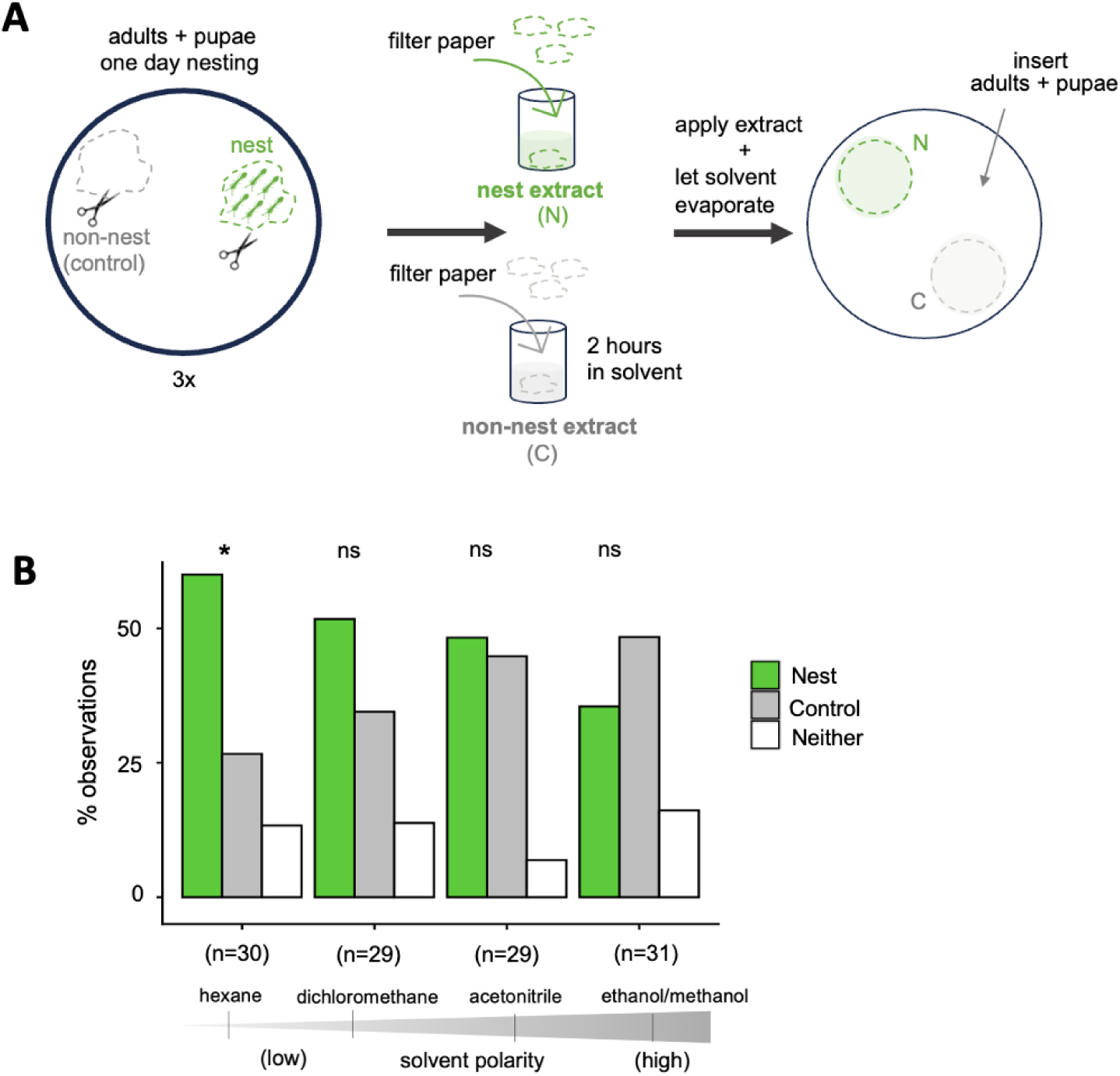
Hexane dissolves an arrestant cue from the nest. **(A)** Experimental design for generating and testing nest and control extracts. Three colonies, each containing 300 adults and 150 pupae, were allowed to establish nests for 1 day. Patches of filter paper were then collected from the nest area and from an area in which the ants had not nested (control). Patches from the three replicate colonies were pooled for extraction. The resulting extracts were applied to new behavioral arenas, after which a separate group of 100 ants (mixed age) in the reproductive phase, together with 50 pupae, was introduced and allowed to form a nest aggregate. **(B)** Percentage of observations of whether colonies nested on the nest extract, the control (non-nest) extract, or neither. Colonies preferentially aggregated on sites treated with hexane extracts from nest patches rather than control patches. A significant preference was absent when more polar solvents were used for extraction. ns, *P* > 0.05; *, *P* < 0.05.

**Figure S3.**
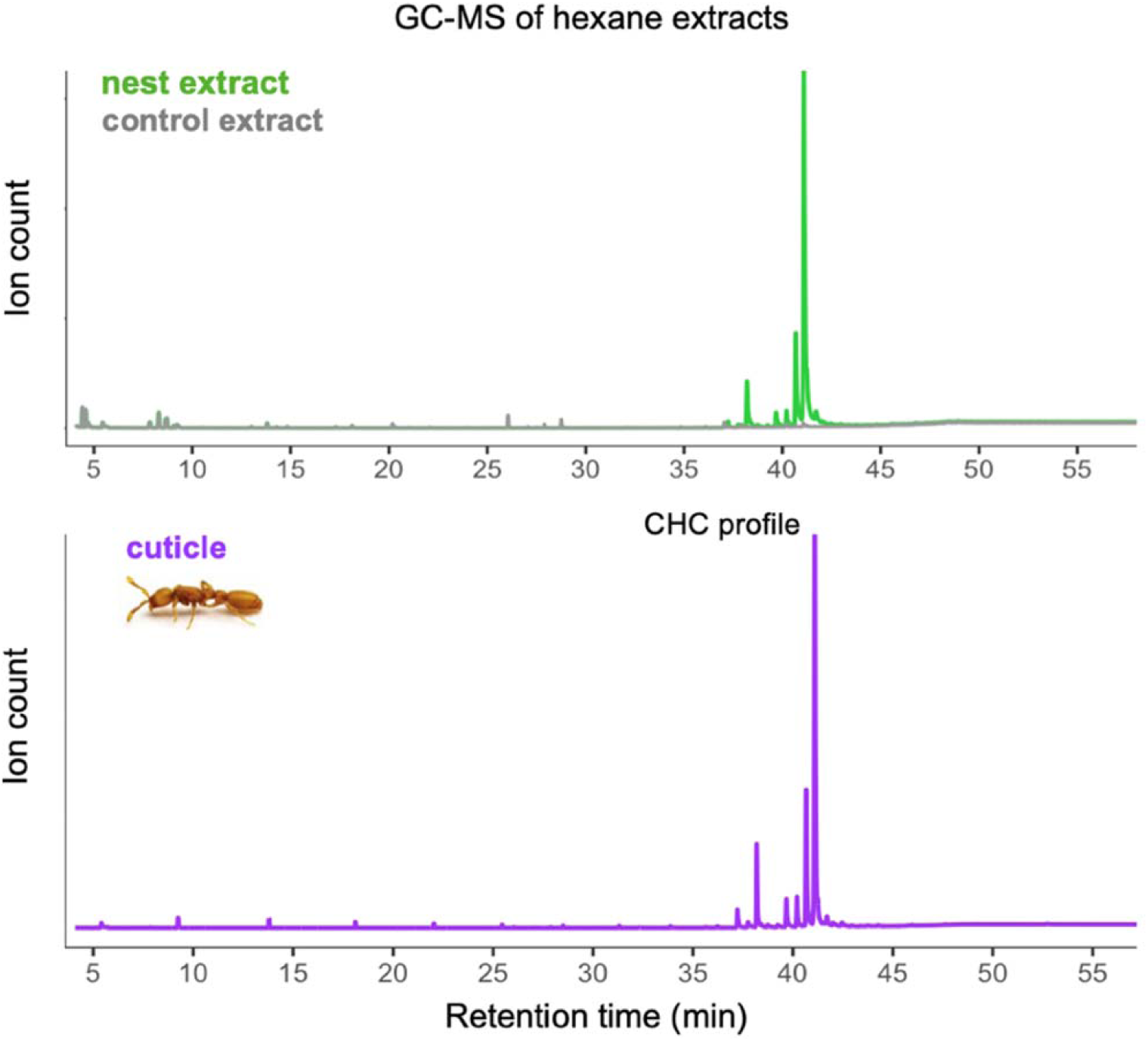
Cuticular hydrocarbons are transferred to the nest. GC–MS profiles of hexane extracts from nest (green) and control (gray) patches of filter paper (top), and of a hexane surface extract from an *O. biroi* worker (purple, bottom). The chemical profile of nest extracts closely matches the ant’s CHC profile.

**Figure S4.**
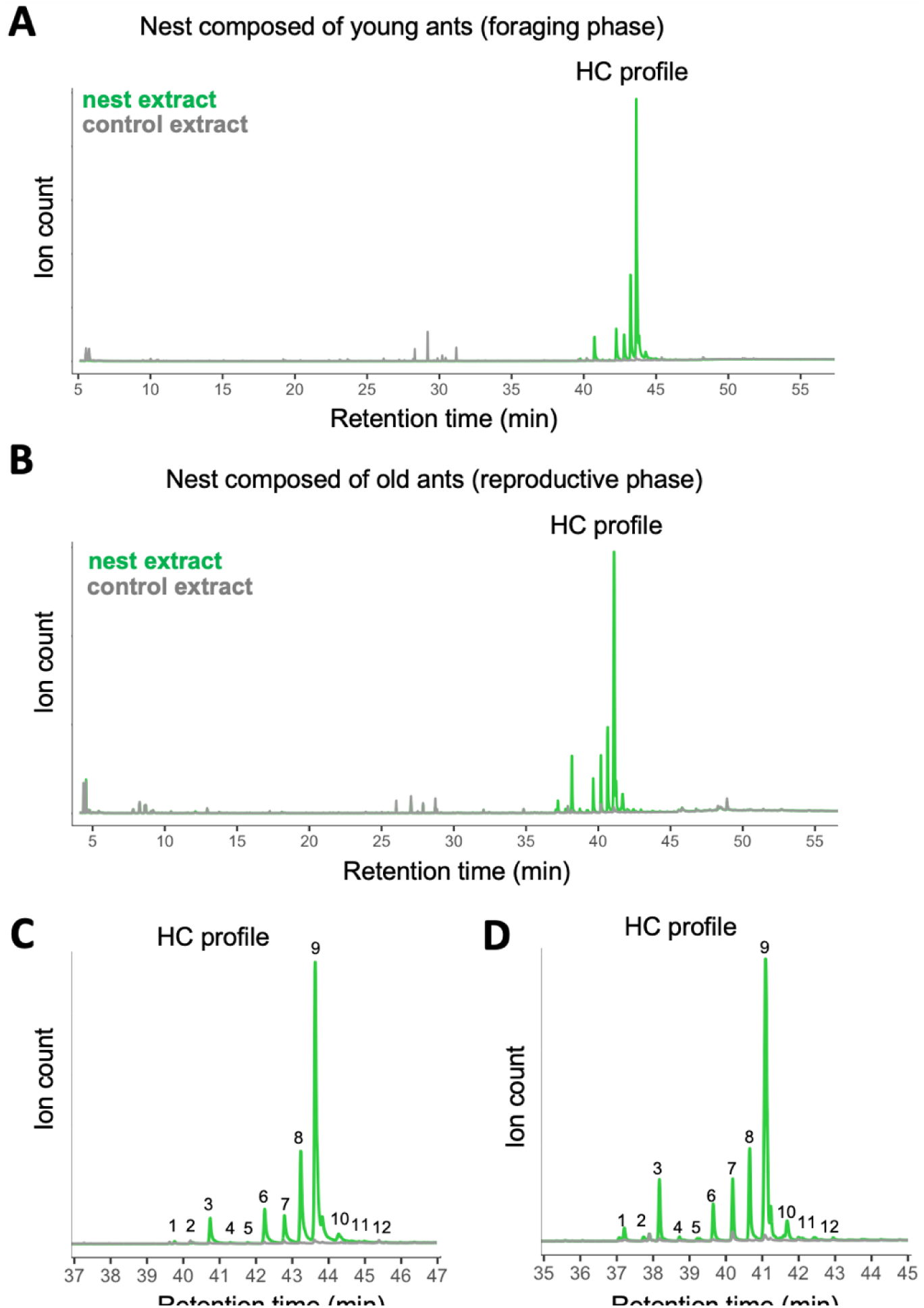
CHCs are transferred to the nest substrate in colonies composed only of adults. GC–MS profiles of nest (green) and non-nest (gray) extracts from colonies composed exclusively of **(A)** young or **(B)** old adult ants. Unlike the colonies shown in Fig. 3A and Fig. S3, these colonies contained no brood. Samples from colonies with young and old ants were analyzed in consecutive years using an identical protocol. Minor shifts in retention time between the two datasets reflect normal instrument drift over this interval. In both age groups, nest extracts contained prominent hydrocarbon (HC) peaks that were greatly reduced in non-nest controls. **(C)** Same as in (A), zooming in the hydrocarbon profile interval (colony of young ants) **(D)**. Same as in (B), zooming in the hydrocarbon profile interval (colony of old ants). HC compounds are listed in the following key: 1 = C25, 2 = 13-;11-MeC25, 3 = 2-;3-MeC25, 4 = C26; 3,7-DiMeC25, 5 = 13-;12-MeC26, 6 = 2-;3-MeC26, 7 = C27, 8 = 13-;11-MeC27, 9 = 11,15-DiMeC27; 2-MeC27, 10 = 3-MeC27, 11 = 3,11-;3,9-;3,7-DiMeC27, 12 = 11,15-DiMeC28; 2-MeC28, (see also Table S1).

**Figure S5.**
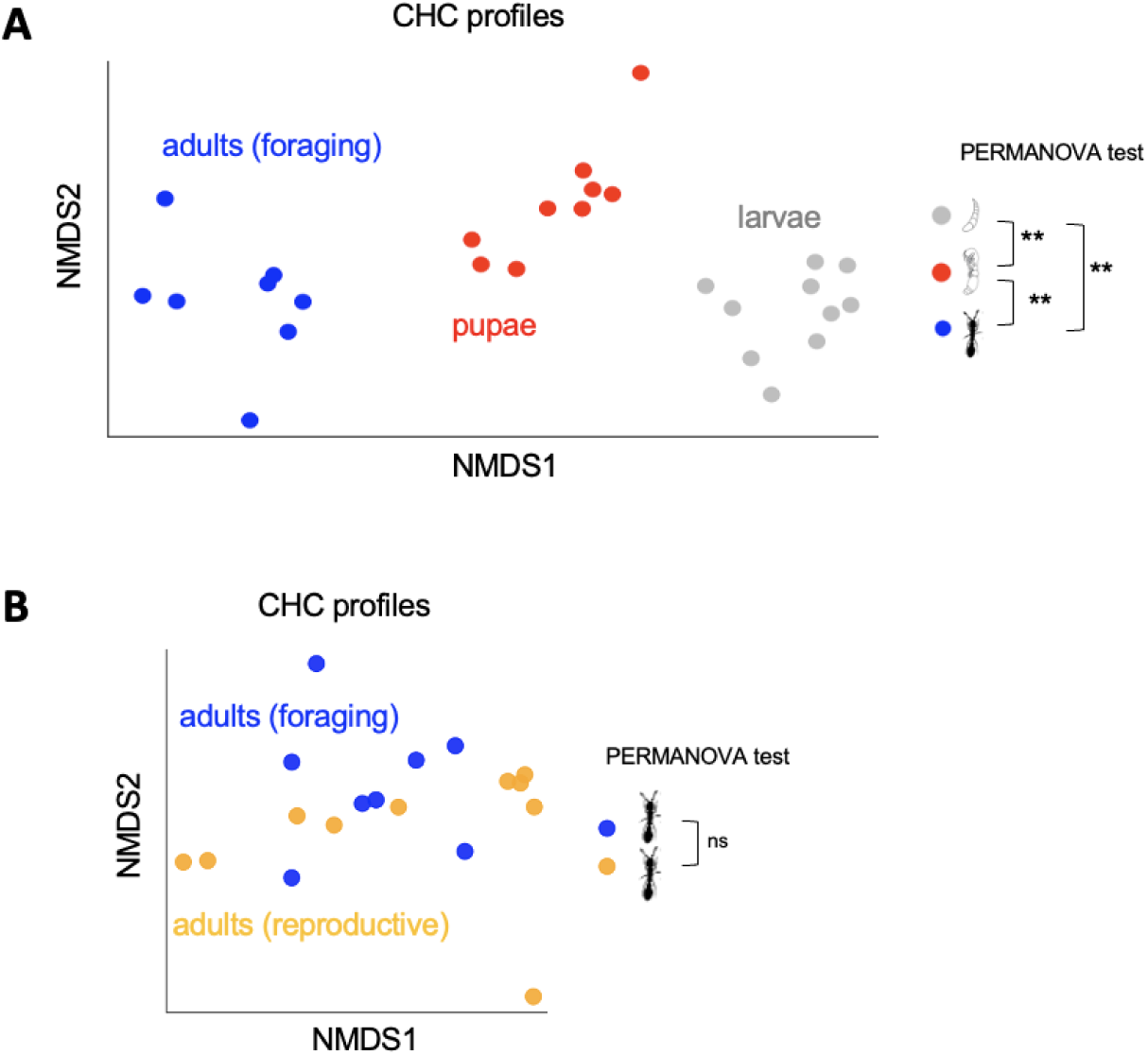
CHC profiles from different developmental stages and adults in different phases of the colony cycle. **(A)** Nonmetric multidimensional scaling (NMDS) analysis of CHC profiles from adults during the foraging phase, as well as larvae and pupae. Profiles clustered by developmental stage and differed significantly between larvae, pupae, and adults. **(B)** NMDS analysis of adult ants during the foraging phase and during the reproductive phase, showing no significant difference. ns, *P* > 0.05; **, *adj. P* < 0.01.

**Figure S6.**
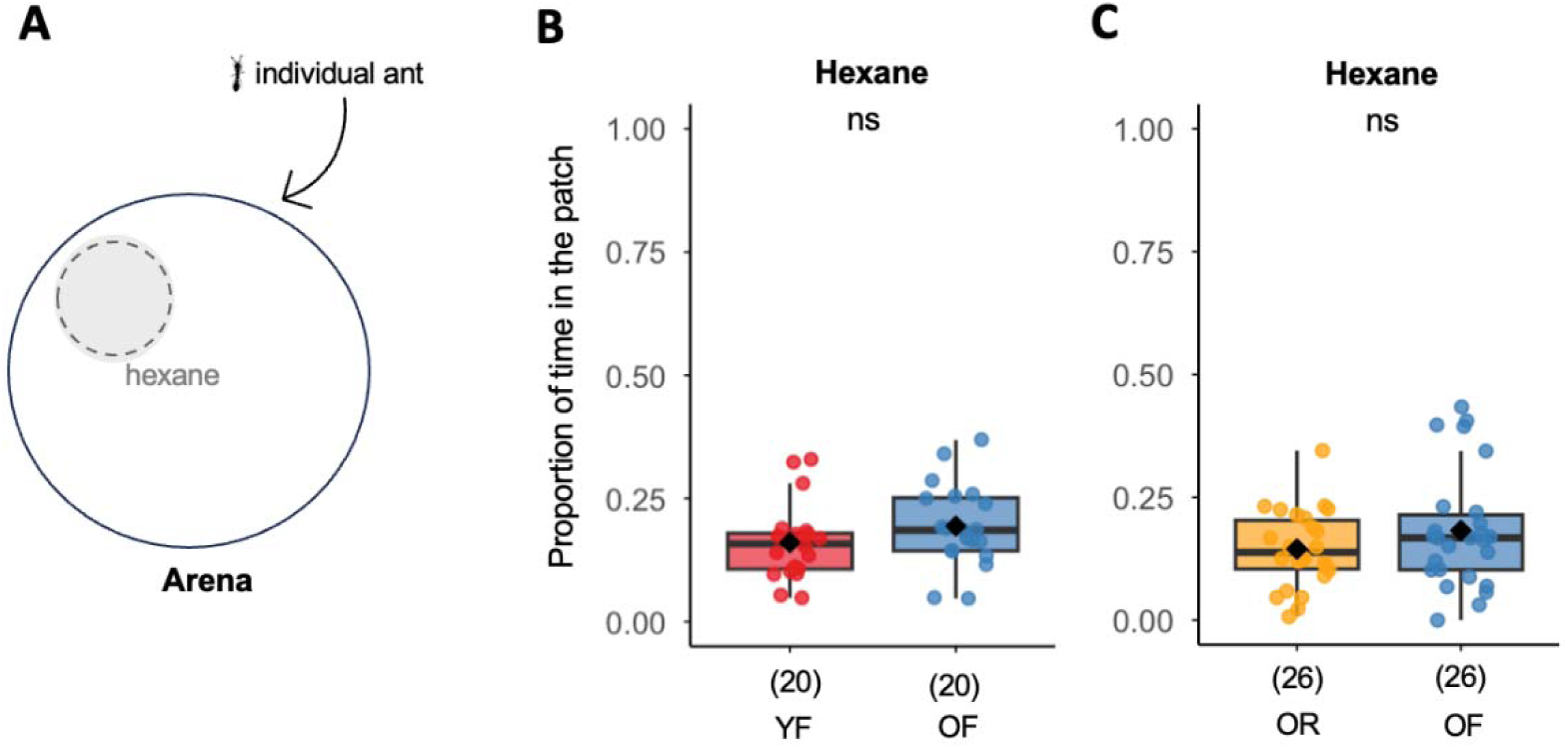
Ants of different ages and from different cycle phases are not differentially arrested by solvent controls. **(A)** Schematic of the experiment. Individual ants were introduced into arenas containing a circular patch treated with hexane. Arrestment was quantified as the proportion of the 30-minute test phase spent within the hexane-treated area. **(B)** Young (5–6 days old) ants (YF, red) and old (4 months old) ants (OF, blue) from colonies in the foraging phase showed no significant difference in time spent on the hexane-treated patch. **(C)** Arrestment did not differ between old ants (>1.7 months old) from colonies in the reproductive phase (OR, orange) and old ants (>1.2 months old) from colonies in the foraging phase (OF, blue). Each dot represents an individual ant. Boxplots delineate the upper and lower quartiles, whiskers represent 1.5 times the interquartile range, diamonds indicate the mean, and horizontal bars the median. Sample sizes are reported on the x-axes. ns, *P* > 0.05.

**Figure S7.**
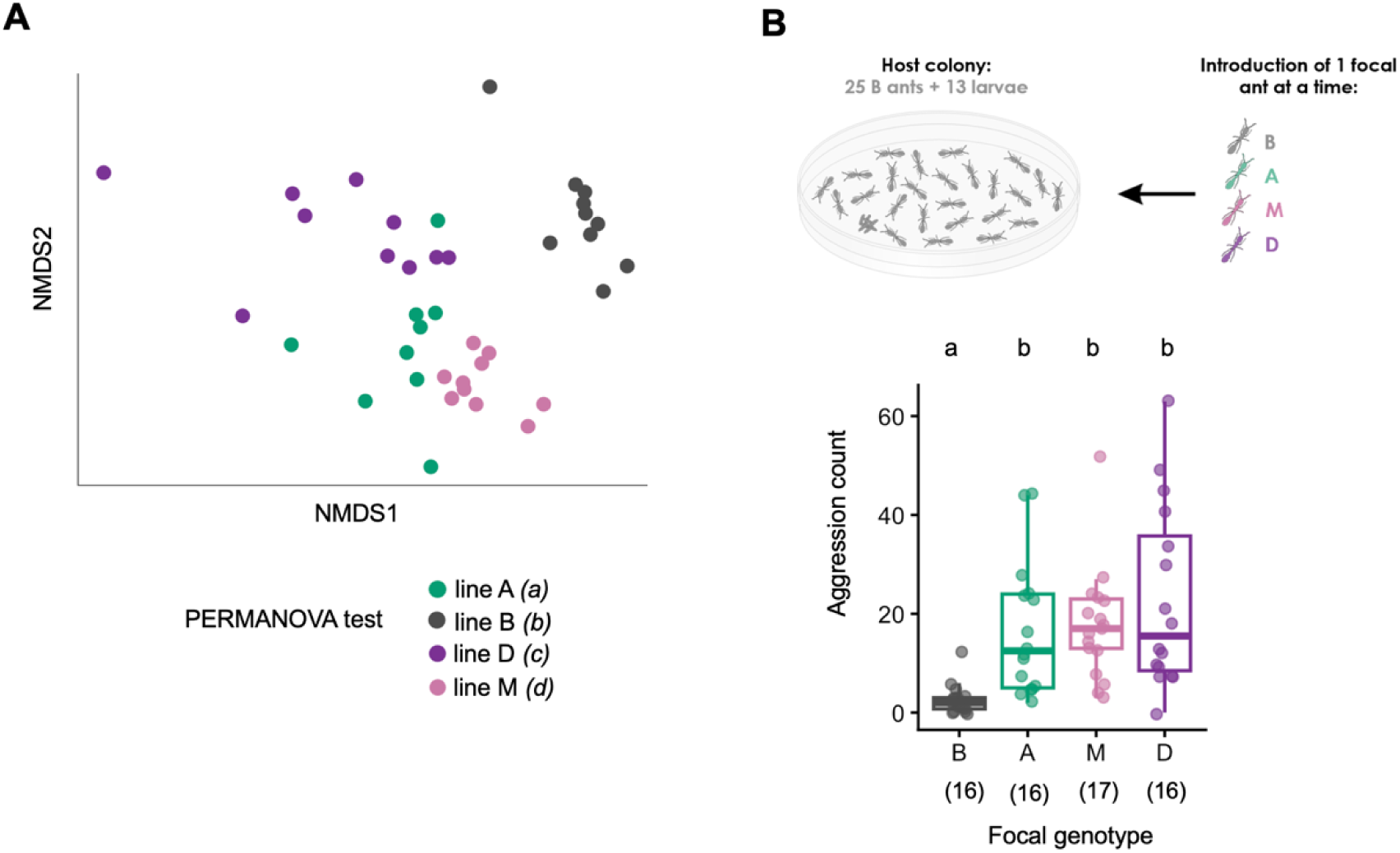
Ants behave aggressively towards adults from other clonal lines during direct encounters. **(A)** Nonmetric multidimensional scaling (NMDS) of CHC profiles from cuticular extracts of adult ants from clonal lines A, B, D and M. Each point represents one CHC profile (a single ant was analyzed for each cuticular extract). Cuticular profiles separated by clonal line in ordination space (PERMANOVA test for overall significant differences between genotypes *P* < 0.001). Different letters in parentheses indicate significant differences between each pair of clonal lines based on pairwise PERMANOVA tests with Bonferroni-Holm correction (all *adj. P* < 0.01). **(B)** Behavioral responses of line B host colonies toward focal ants from different clonal lines. Individual focal ants from clonal lines B, A, M and D were sequentially introduced into host colonies of 25 line B adults and 13 larvae in randomized order. Aggression counts represent the total number of aggressive acts directed by host workers toward each focal ant. Line B ants from a different colony elicited significantly less aggression than focal ants from the three other clonal lines, whereas aggression levels towards ants from lines A, M, and D did not differ significantly. Each dot represents one focal ant, boxes show the interquartile range, whiskers represent 1.5 times the interquartile range, and horizontal lines indicate medians. Different letters indicate significant differences (*P* < 0.001) between groups based on a generalized linear mixed model with a negative binomial distribution and date of the experiment as a random factor (as in^1^). Data for focal ants from lines A, B, and D were previously reported^1^, along with a detailed description of the aggression assay methods. Data for line M are reported here for the first time but were obtained concurrently with the data for the other clonal lines.

**Figure S8.**
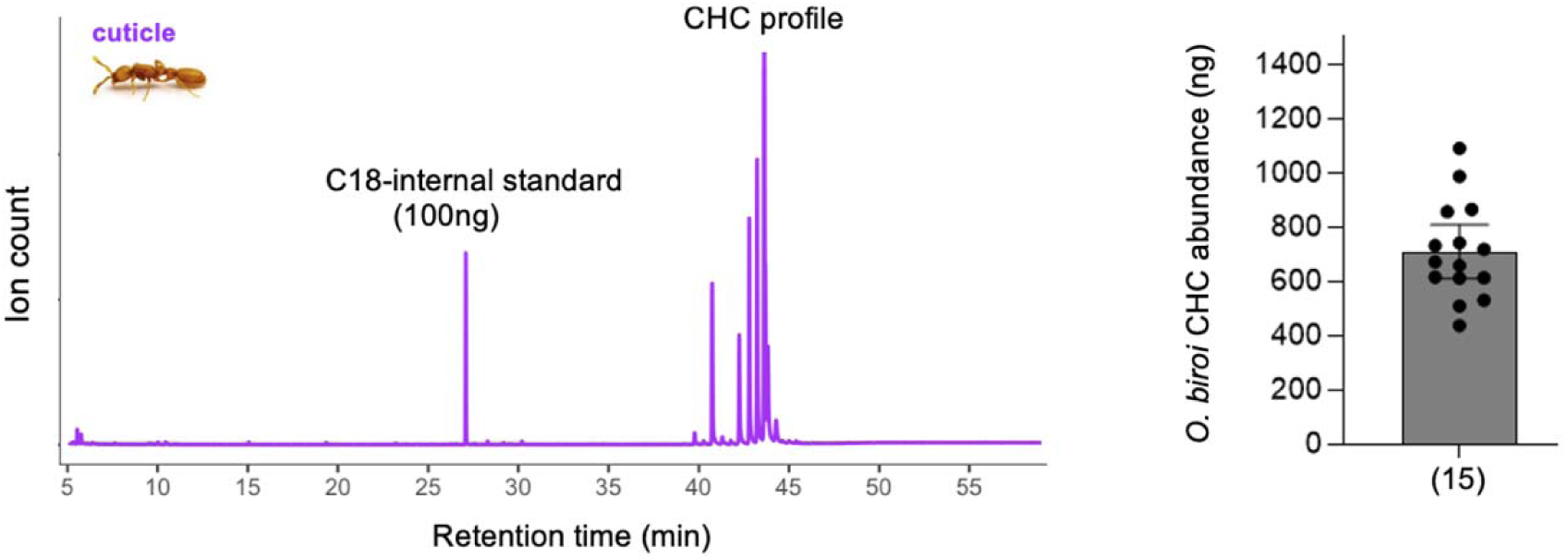
Quantification of *O. biroi* CHCs. Left: Chromatogram of hexane extract from an adult *O. biroi* ant, showing the internal C18 standard (100 ng) used for quantification. Right: Total CHC abundance (mass) for n = 15 individual age-matched (2-weeks old) *O. biroi* ants. The bar shows the mean, and whiskers indicate ± the standard error of the mean.

**Table S1.** CHC compounds identified across *O. biroi* clonal lines. The table lists the retention index (RI) and the relative abundance (mean ± standard deviation) for each CHC compound that was identified in hexane extracts from adult ants from different clonal lines and in hexane extracts from nest substrates of clonal line B colonies. Semicolons separate compounds that were detected but could not be reliably separated in all samples. Asterisks indicate compounds that could not be separated due to peak tailing.

| **Compound** | **RI** | **cuticle A** | **cuticle B** | **cuticle D** | **cuticle M** | **nest B** |
|---|---|---|---|---|---|---|
| C25 | 2500 | 0.65 ± 0.18 | 1.15 ± 0.24 | 0.36 ± 0.09 | 0.80 ± 0.18 | 2.59 ± 0.48 |
| 13-;11-MeC25 | 2533 | 0.54 ± 0.16 | 0.95 ± 0.17 | 0.71 ± 0.23 | 0.91 ± 0.05 | 1.20 ± 0.44 |
| 2-;3-MeC25* | 2564 | 9.38 ± 1.31 | 11.22 ± 0.95 | 8.50 ± 0.61 | 6.44 ± 0.49 | 11.05 ± 0.64 |
| C26; 3,7-DiMeC25 | 2600 | 0.83 ± 0.13 | 0.62 ± 0.14 | 0.56 ± 0.22 | 0.94 ± 0.11 | 0.58 ± 0.18 |
| 13-;12-MeC26 | 2632 | 0.95 ± 0.22 | 0.97 ± 0.18 | 0.81 ± 0.31 | 1.05 ± 0.15 | 1.05 ± 0.37 |
| 2-;3-MeC26* | 2663 | 6.09 ± 0.58 | 5.06 ± 0.45 | 4.80 ± 0.48 | 7.49 ± 0.64 | 5.45 ± 1.12 |
| C27 | 2700 | 6.04 ± 1.28 | 4.57 ± 1.18 | 3.41 ± 0.60 | 5.13 ± 1.15 | 5.53 ± 1.45 |
| 13-;11-MeC27 | 2730 | 14.88 ± 1.35 | 18.75 ± 0.80 | 17.98 ± 0.55 | 15.63 ± 0.88 | 19.00 ± 1.23 |
| 11,15-DiMeC27; 2-MeC27 | 2757 | 45.58 ± 2.51 | 48.23 ± 3.24 | 46.46 ± 2.27 | 46.95 ± 1.05 | 44.87 ± 4.35 |
| 3-MeC27 | 2771 | 6.84 ± 0.91 | 4.11 ± 0.55 | 7.39 ± 0.97 | 6.55 ± 0.45 | 5.72 ± 1.01 |
| 3,11-;3,9-;3,7-DiMeC27 | 2801 | 7.51 ± 0.99 | 3.57 ± 0.75 | 8.36 ± 1.41 | 7.51 ± 1.12 | 2.82 ± 1.06 |
| 11,15-DiMeC28; 2-MeC28 | 2854 | 0.70 ± 0.26 | 0.78 ± 0.13 | 0.67 ± 0.28 | 0.58 ± 0.22 | 0.14 ± 0.11 |

### Appendix

The syntheses of the following compounds have been previously published: 3Me-C27^7^, 11Me-C27^8^, 13Me-C27^9^, 2Me-C30^10^. The remaining compounds were synthesized as described below. Tetrahydrofuran (THF) was dried by distillation from benzophenone ketyl under Ar. Unless otherwise specified, solutions were dried with anhydrous Na_2_SO_4_ and concentrated by rotary evaporation. Vacuum flash chromatography was performed with 230-400 mesh silica gel in sintered glass funnels. Air- and water-sensitive reactions were run in oven-dried glassware under Ar. The identities of intermediates were checked by coupled gas chromatography-mass spectrometry, using a Hewlett-Packard (now Agilent, Santa Clara CA, USA) 6890 GC interfaced to a 5973-mass selective detector, with electron impact ionization (70 eV). The GC was equipped with a DB-17 column (30 m x 0.25 mm id, J&W Scientific, Folsom CA, USA). The GC was typically programmed from 40°C/1 min, 10°C /min to 280°C, hold 20 min.

#### Synthesis of 2-methylpentacosane

Undecanol (0.93 g, 5 mmol), pyridine (0.47 g, 6 mmol), and dimethylaminopyridine (50 mg) were added to 15 ml CH_2_Cl_2_, and the solution was cooled in an ice bath while adding triflic anhydride (1.69 g, 6 mmol) over 30 min with a syringe pump. The mixture was stirred at 0°C for 1 h, then warmed to room temperature and stirred 1 h. The mixture was diluted with 30 ml hexane, stirred 5 min, then filtered through a pad of celite, rinsing with 50 ml 1:1 hexane:CH_2_Cl_2_. After concentration, the pink residue was pumped briefly under vacuum, then taken up in 5 ml ether and used immediately.

An oven-dried 3-neck flask was flushed with Ar and charged with 13-methyltetradec-1-yne (0.83 g, 4 mmol), ∼25 mg triphenylmethane indicator, and 20 ml dry THF. The solution was cooled in an ice bath, and butyllithium (2.5 M in hexanes) was added dropwise until the solution turned pink, followed by an additional 1.6 ml BuLi (4 mmol). The solution was stirred 30 min, then the ethereal solution of the triflate was added by syringe pump over 30 min. The mixture was warmed to room temperature and stirred 1 h, then quenched with saturated aqueous NH_4_Cl, and extracted with hexane. The hexane solution was washed with brine, dried, and concentrated, and purified by vacuum flash chromatography, eluting with hexane. The fractions containing the desired alkyne product were combined and concentrated, and the residue was taken up in 15 ml hexane. After addition of 50 mg 10% Pd on carbon catalyst, the mixture was flushed with Ar and then hydrogenated with a balloon of hydrogen for 2 h. The mixture was then filtered through celite, and the residue was Kugelrohr distilled (100°C, 0.05 mm Hg) to remove traces of 13−methyltetradecane. The pot residue was then recrystallized from 15 ml acetone, chilling in a −20°C freezer overnight. After filtration and air drying, 2-methylpentacosane was recovered as fine white crystals (0.48 g, 33%). The synthesis scheme and the EI mass spectrum (70 eV) are shown below.

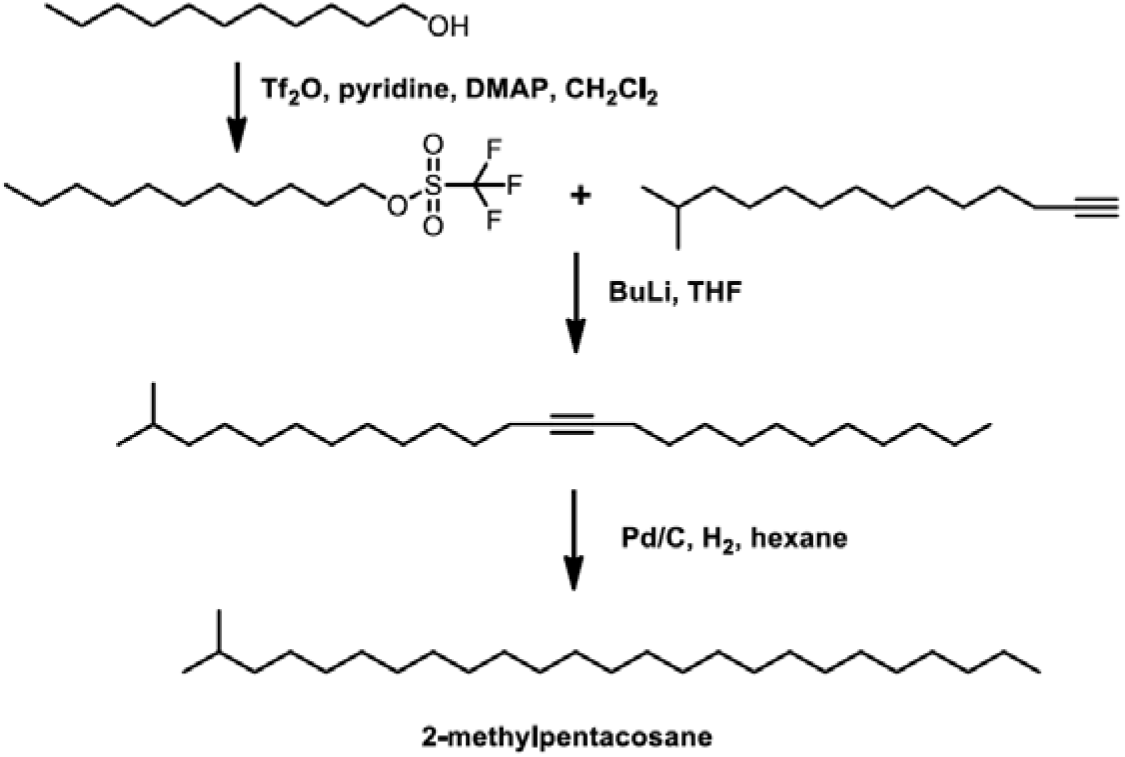

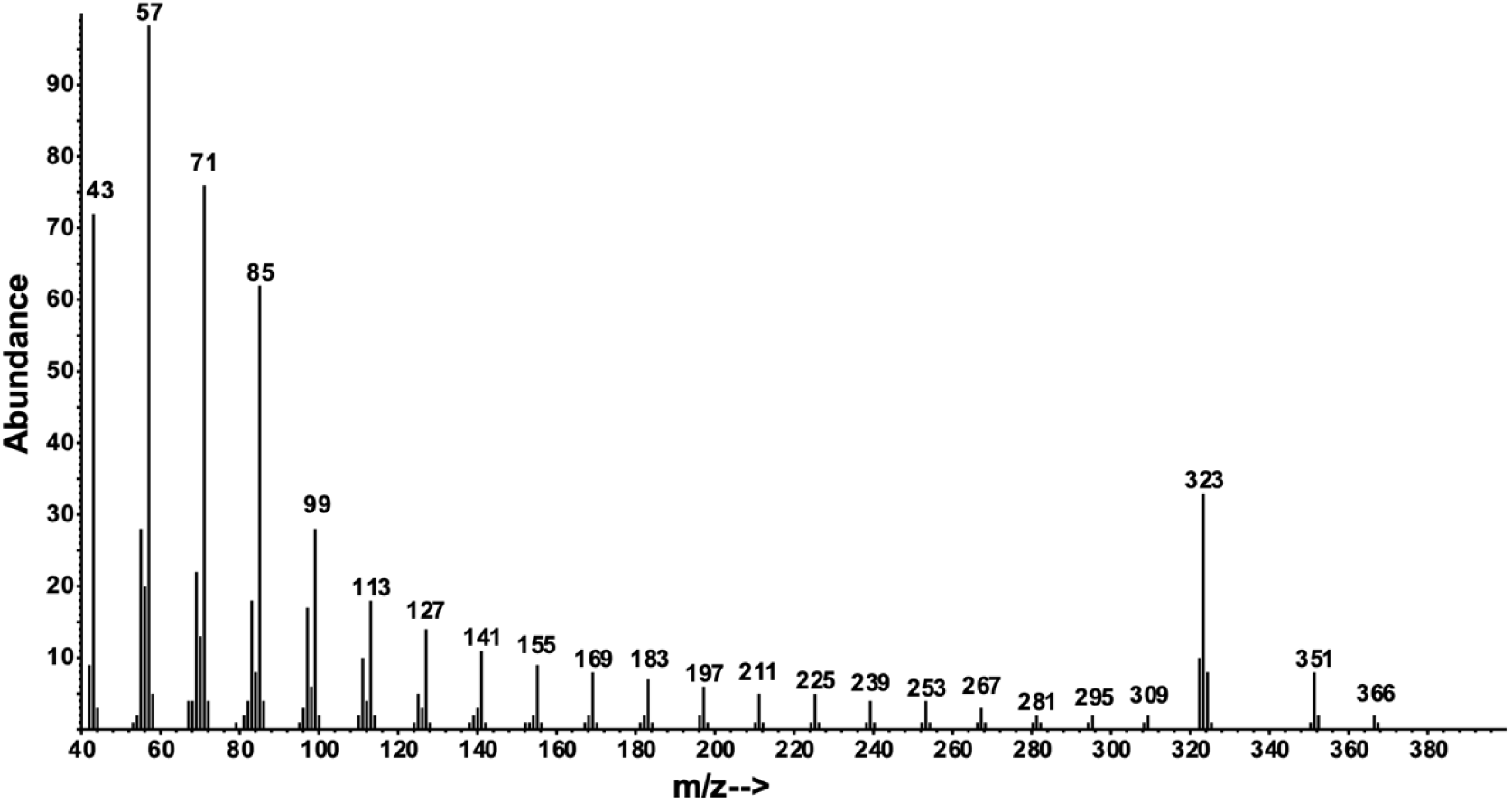

#### Synthesis of 3,9-dimethylheptacosane

2-Methyleicosanol: The triflate of octadecanol was prepared as described above for the analogous undecyl triflate, taken up in ether, and used immediately as described below.

2-Ethyl-4,4-dimethyl-2-oxazoline (3.17 g, 25 mmol) was dissolved in 20 ml dry THF under argon, the solution was cooled to −78°C, BuLi (2.5 M in hexanes, 10 ml, 25 mmol) was added dropwise over 30 min, and the mixture was stirred an additional 30 min at −78°C. The ethereal solution of the crude triflate (nominal, 25 mmol) was then added in four aliquots over 30 min, resulting in a yellow slurry, which was allowed to warm to room temperature overnight. The mixture was quenched with saturated brine and extracted with ether, which after concentration yielded 6.4 g of a viscous yellow oil which started to crystallize. Aqueous 3M HCl (40 ml, 120 mmol) was added, and the mixture was refluxed for 45 min, yielding a yellow slurry. The mixture was cooled, diluted with water, and extracted twice with CH_2_Cl_2_. The crude product was subjected to an acid-base extraction by taking it up in 0.5 M NaOH (100 ml) and extracting twice with ether. The aqueous solution was then acidified with 20 ml 4 M HCl and extracted twice with ether. The ethereal solution precipitated some off-white solids, and so was filtered, washed with brine, and concentrated, yielding 1.7 g of an off-white solid. This was taken up in 25 ml EtOH with 1 ml conc. sulfuric acid, and the mixture was refluxed for 2 h. The mixture was then concentrated, partitioned between water and ether, and the ethereal solution was washed with saturated aqueous NaHCO_3_ and brine, then concentrated and purified by vacuum flash chromatography (10% EtOAc in hexane), yielding ethyl 2-methyleicosanoate (1.32 g, 15%). EIMS (70eV; *m/z*, %): 354 (M^+^,26), 311 (5), 297 (9), 281 (2), 269 (2), 255 (3), 241 (3), 227 (3), 213 (3), 199 (2), 185 (1), 171 (10), 157 (9), 129 (4), 115 (37), 102 (100), 87 (9), 74 (13), 69 12(), 57 17(), 55 15(), 43 (19), 41 (11).

The ester (1.25 g, 3.5 mmol) in 5 ml THF was added dropwise to a slurry of LiAlH_4_ (0.19 g, 5 mmol) in 10 ml THF cooled in an ice bath, under Ar. When the addition was complete, the mixture was warmed to room temperature and stirred 3 h, then cooled again, and 0.2 ml water, 0.17 ml 20% aqueous NaOH, and 0.7 ml water were added dropwise, sequentially, over 20 min. The resulting slurry was warmed to room temperature and stirred 30 min, then filtered through a pad of celite, rinsing well with ether. The filtrate was dried, then concentrated to 2-methyleicosanol (white solid, 0.98 g, 90%, 98% pure by GC), which was used without further purification. EIMS (70 eV; *m/z*, %): 294 (M^+^ −18,7), 266 (8), 252 (2), 238 (3), 224 2(2), 210 (2), 196 (2), 182 (3), 168 (4), 153 (6), 139 (12), 125 (26), 111 (48), 97 (80), 85 (44), 83 (69), 71 (61), 69 (69), 57 (100), 56 47(), 55 (68), 43 (77), 41 (51).

5-Methylheptyl bromide: ten ml of a solution of 1-bromo-2-methylbutane (5 g, 33 mmol) in THF (60 ml) was added in one portion to a dry flask under Ar, charged with Mg chips (1.2 g, 50 mmol), and the mixture was stirred until it warmed and turned grey, at which point the remaining solution was added dropwise over 1 h. The mixture was then warmed to 45°C for 1 h, then cooled to room temperature.

A separate dry flask under Ar was charged with CuI (0.38 g, 2 mmol) and 20 ml THF, then cooled to ∼-15°C in an ice-acetone bath, and the Grignard reagent prepared above was added dropwise over 1 h, followed by addition of β-propiolactone (1.8 g, 25 mmol) in 5 ml THF over 30 min by syringe pump. The mixture was allowed to warm to room temperature and stirred overnight, then quenched with 100 ml 1M H_2_SO_4_. The mixture was extracted twice with ether, and the combined ether extracts were back-extracted with 0.5 M KOH (75 ml). The aqueous layer was then acidified with 6 M H_2_SO_4_ and extracted 3 times with ether. The combined ether extracts were washed with brine, dried, and concentrated, yielding 5-methylheptanoic acid as a yellow oil (3.39 g, 94%).

The acid (3.2 g, 22.2 mmol) was taken up in 5 ml ether, and added dropwise to an ice-bath cooled slurry of LiAlH_4_ (0.95 g, 25 mmol) in ether (50 ml) under Ar. The resulting mixture was warmed to room temperature and stirred 3 h, then cooled to 0°C again and quenched by successive dropwise addition of 1 ml water, 0.75 ml 20% aqueous NaOH, and 3.5 ml water. The resulting slurry was stirred 30 min, then filtered, rinsing well with ether. The filtrate was dried and concentrated, yielding 5-methylheptanol (1.9 g, 66%). EIMS (70eV; *m/z*, %): 101 (M^+^-29,1), 97 (7), 84 (22), 83 (86), 82 (12), 73 (10), 70 (46), 69 (21), 57 (29), 56 (41), 55 (100), 43 (20), 42 (10).

The alcohol (1.8 g, 13.8 mmol) and triphenylphosphine (4.98 g, 19 mmol) were dissolved in 20 ml CH_2_Cl_2_, the solution was cooled in an ice bath, *N*-bromosuccinimide (2.49 g, 18 mmol) was added in portions over 30 min, and the mixture was warmed to room temperature for 3 h. The mixture was then concentrated, and the residue was triturated with 50 ml pentane. The pentane solution was filtered through a plug of silica gel, rinsing with pentane, and the filtrate was concentrated, yielding 1.48 g (56%) of 5-methylheptyl bromide as a colorless liquid. EIMS (70 eV; *m/z*, %): 165, 163 (M^+^-29, 8), 137 (97), 135 (100), 123 (5), 121 (5), 109 (3), 107 (3), 97 (5), 83 (60), 71 (5), 69 (7), 57 (67), 56 (30), 55 (75), 43 (27).

3,9-dimethylheptacosane: 2-methyleicosyl triflate was prepared from 2-methyleicosanol as described above, taken up in 5 ml ether, and used immediately as described below. 5-Methylheptylmagnesium bromide was prepared from 5-methylheptyl bromide and Mg chips in THF as described above, and the grey solution was added dropwise over 30 min to a slurry of CuI (76 mg, 0.4 mmol) in 10 ml THF under Ar, cooled to ∼-15°C in an ice-salt bath, giving a pale grey slurry, which was stirred 30 min, then warmed to ∼-10°C. The ethereal solution of 2-methyleicosanyl triflate was then added by syringe pump over 30 min, and the mixture was allowed to warm to room temperature overnight. The mixture was then quenched with saturated NH_4_Cl and extracted with ether. After concentration, the crude product was found to be heavily contaminated with 2-methyleicosanyl bromide and iodide, and these contaminants were removed as follows.

A dry flask under Ar was charged with NaH (60% dispersion in oil, 1 g, 25 mmol), which was rinsed twice with hexane. THF (25 ml) and DMF (6 ml) were then added, followed by dropwise addition of diethylmalonate (4 g, 22.5 mmol) over 1 h. The resulting slurry was stirred 1 h, then NaI (0.75 g, 5 mmol) was added, followed by dropwise addition of a solution of the impure 3,9-dimethylheptacosane in 5 ml THF over 30 min. The mixture was warmed to 45°C and stirred overnight, then cooled to room temperature, quenched with saturated aqueous NH_4_Cl, and extracted with hexane. The hexane extract was washed with water and brine, dried, and concentrated. The crude product was Kugelrohr distilled (oven temperature to 120°C at 0.1 mm Hg) to remove excess diethylmalonate. The pot residue was then purified by vacuum flash chromatography, eluting with hexane, with a final purification by recrystallization twice from hot acetone, yielding the desired product (120 mg) in >98% purity by GC. The synthesis scheme and the EI mass spectrum (70 eV) are shown below.

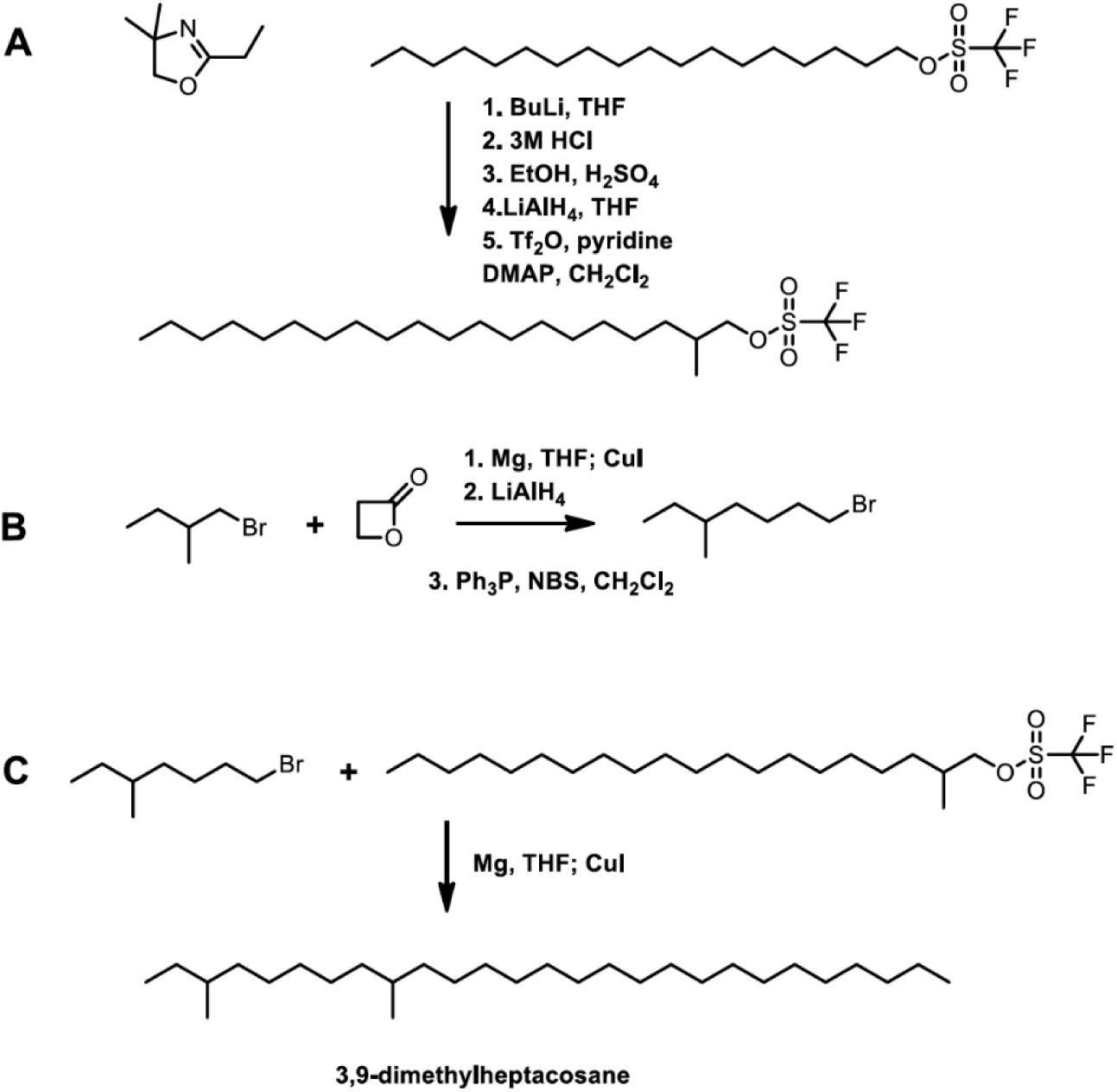

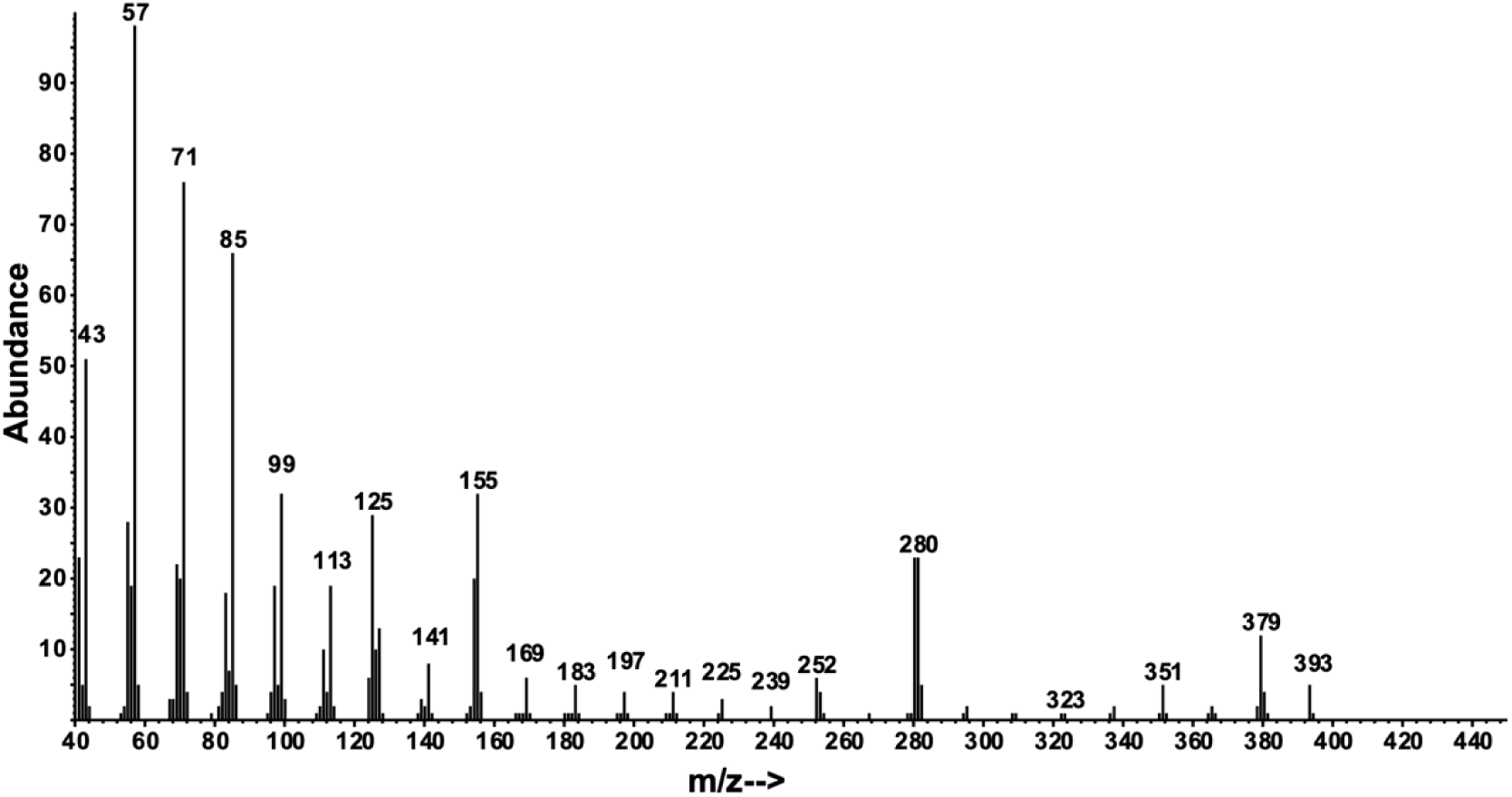

#### Synthesis of 11,15-dimethylheptacosane

6-Hydroxy-2-methylheptanoate: a 500 ml flask was charged with 6 M H_2_SO_4_ (80 ml), cooled in an ice-bath, and ammonium peroxysulfate (18.24 g, 80 mmol) was added in portions over 5 min, followed by dropwise addition of 2,6-dimethylcyclohexanone (5.04 g, 40 mmol). The mixture was stirred 30 min, then the resulting clear solution was extracted with EtOAc. The EtOAc solution was washed with brine and dried, and contrary to the literature report^11^, appeared to be almost exclusively 6-hydroxy-2-methylheptanoic acid, rather than the corresponding lactone. Thus, the crude hydroxyacid was added to a solution of HCl in MeOH, generated by dropwise addition of 1.5 ml conc. HCl to 50 ml MeOH at 0°C, and the solution was heated to 60°C for 1 h. The mixture was allowed to cool to room temperature overnight, then diluted with 150 ml water and extracted 3 times with ether. The combined ether extracts were washed with saturated aq. NaHCO_3_, then brine, and dried. The solution was concentrated and the residue was purified by Kugelrohr distillation (bp∼80°C, 1 mm Hg), yielding methyl 6-hydroxy-2-methylheptanoate (4.02 g, 58%) as a colorless oil. EIMS (70 eV; *m/z*, %): 159 (M^+^-15,2), 143 (5), 130 (31), 127 (20), 125 (11), 115 (4), 102 (6), 101 (84), 99 10(), 98 (10), 97 (23), 88 (76), 87 (100), 74 4(), 73 (10), 71 (8), 70 (6), 69 (24), 59 (24), 57 (23), 56 (11), 55 (49), 45 (47), 43 (32), 42 (14), 41 (27).

Methyl 2-methyl-6-oxo-heptanoate: Methyl 6-hydroxy-2-methylheptanoate (1.74 g, 10 mmol) was added dropwise over 30 min to a solution of Dess-Martin periodinane (5.1 g, 12 mmol) in 100 ml CH_2_Cl_2_, and the mixture was stirred at room temperature for 3 h. The mixture was then concentrated by rotary evaporation, the residue was taken up in 100 ml ether and 150 ml 10% aqueous sodium thiosulphate, and the mixture was stirred vigorously for 1 h. The layers were separated, and the ether layer was washed with water and brine, dried, and concentrated. The residue was triturated with hexane, and the hexane soluble portion was purified by vacuum flash chromatography on silica gel, eluting with 25% EtOAc in hexane. The purified material was then Kugelrohr distilled (bp∼90°C at 3 mm Hg), yielding the ketoester as a colorless oil (1.31 g, 76%). Methyl (*E*/*Z*6)-2,6-dimethylhexadecenoate: Butyllithium (2.5 M in hexanes) was added dropwise to a suspension of decyltriphenylphosphonium bromide (2.9 g, 6 mmol) in 30 ml dry THF under Ar until the mixture turned pale orange, followed by addition of a further 2.4 ml (6 mmol) of BuLi solution. The orange solution was stirred 30 min, then the ketoester (0.86 g, 5 mmol) in 2 ml THF was added dropwise by syringe pump over 30 min. The mixture was stirred overnight, then quenched with saturated aq. NH_4_Cl, and extracted with hexane. The hexane layer was washed with brine, dried, and concentrated. The residue was triturated with hexane, and the hexane soluble portion was purified by vacuum flash chromatography on silica gel, eluting with 4% EtOAc is hexane, yielding the alkenyl ester (400 mg, 23%) as a ∼ 1:1 mixture of (*Z*)- and (*E*)-isomers. (*E*/*Z*6)-2,6-Dimethylhexadecenal: The ester in 5 ml THF was added dropwise to a slurry of LiAlH_4_ in 10 ml THF at 0°C, and the mixture was warmed to room temperature and stirred 1 h. The reaction was then quenched by sequential dropwise addition of 100 µl water, 67 µl 20% aq. NaOH, and 360 µl water. The mixture was stirred for 30 min, then filtered and concentrated. The crude alcohol (as a ∼1:1 mixture of isomers) was taken up in 5 ml CH_2_Cl_2_ and added dropwise to a slurry of Dess-Martin periodinane in 15 ml CH_2_Cl_2_. The mixture was stirred 3 h at room temperature, then concentrated. The residue was taken up in 50 mL ether and stirred vigorously for 20 min with a 1:1 aq. solution of 10% sodium thiosulphate and saturated NaHCO_3_ (60 mL).

The ether layer was then separated, washed with brine, dried, concentrated, and used immediately in the next step. 11,15-Dimethylheptacosane: BuLi (2.5 M in hexanes) was added dropwise to a slurry of undecyltriphenylphosphonium bromide (0.994 g, 2 mmol) in 20 mL dry THF until an orange color persisted, then an additional 0.8 mL BuLi (2 mmol) were added. The resulting orange solution was stirred 1 h, then the crude aldehyde from above in 2 mL THF was added over 30 min with a syringe pump. The solution was stirred at room temperature overnight, then quenched with dilute aq. NH_4_Cl, and extracted with hexane. After concentration, the residue was taken up in ∼5 mL hexane, and purified by vacuum flash chromatography, eluting with hexane. The hydrocarbon fraction (a 100:14:88:19 mixture of diene isomers) was concentrated to ∼25 mL, 50 mg 10% Pd on C catalyst was added, and the mixture was stirred 2 h under H_2_ atmosphere, supplied with a balloon. The mixture was then filtered and concentrated. The product was found to be contaminated with about 10% docosane. Thus, the residue was taken up in 50 mL isooctane, 2 g of freshly dried 5 Å powdered molecular sieve were added, and the slurry was stirred 2 d, then filtered through a plug of cellite, rinsing with isooctane. After concentration, the residue was taken up in 15 mL hot acetone, and recrystallized overnight at ∼-20°C. The mixture was filtered in a cold room, and the resulting white crystals (350 mg, 98% pure) melted at ∼room temperature to a colorless oil. The synthesis scheme and the EI mass spectrum (70 eV) are shown below.

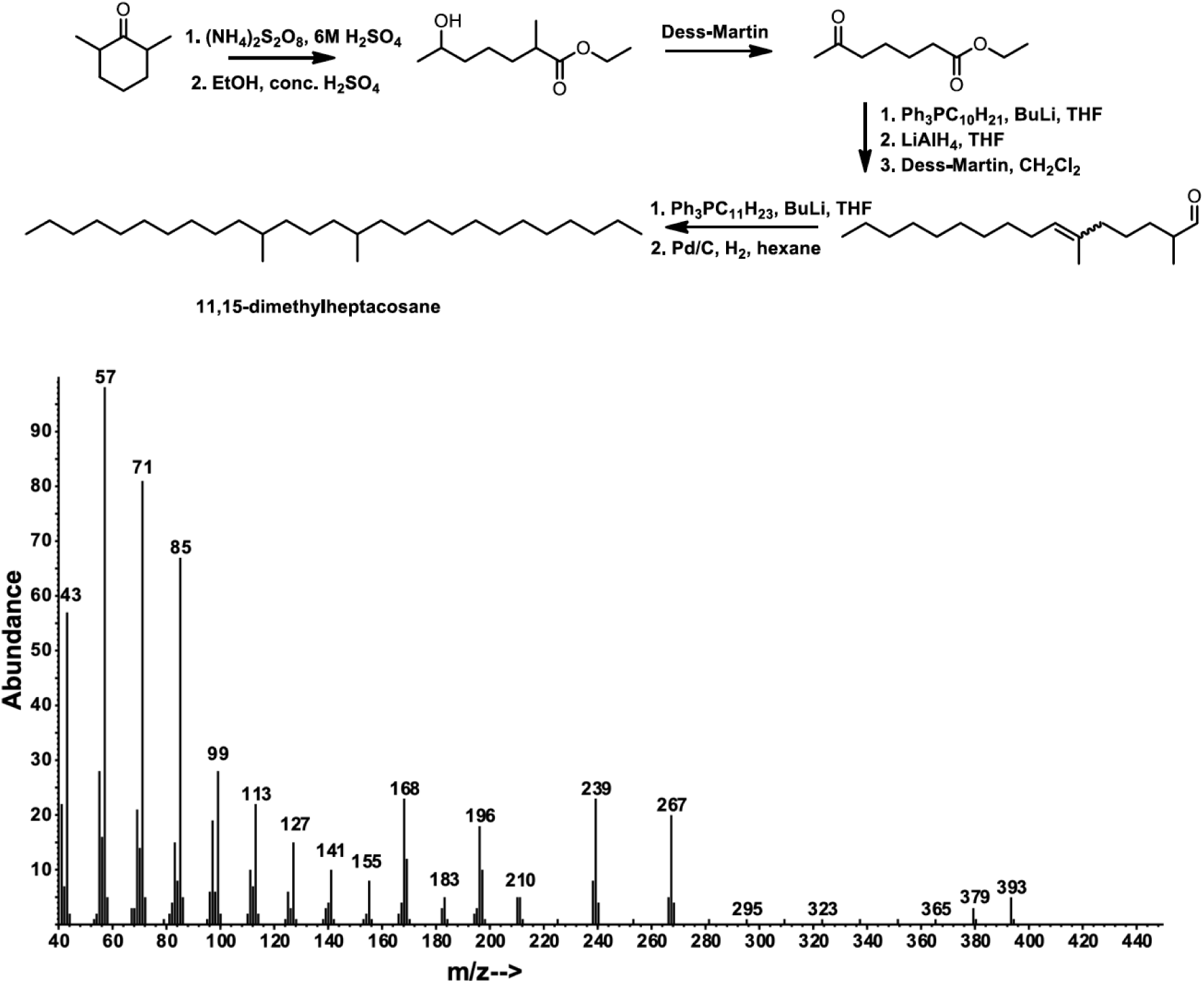

#### Synthesis of 13,23-dimethylhexatriacontane

2-methyltetracosanoic acid: A dry flask under Ar was charged with 75 ml THF and dry diisopropylamine (9.6 ml, 68 mmol), and cooled to −78°C. BuLi (2.5 M, 27.2 ml, 68 mmol) was added dropwise over 30 min, and the mixture was warmed to 0°C and stirred 30 min. A solution of propanoic acid (2.22 g, 30 mmol) in 5 ml THF was then added dropwise, giving a slightly cloudy pale-yellow solution. The mixture was stirred 20 min at 0°C, then N,N′-dimethylpropyleneurea (DMPU, 4.1 ml, 34 mmol) was added, yielding a pale-yellow slurry. The mixture was warmed to room temperature and stirred 30 min, then cooled to 0°C again, and dodecyl bromide (8.7 g, 35 mmol) was added in one portion. The mixture was warmed to room temperature overnight, then warmed to 35°C for 3 h. The mixture was then quenched and acidified with cold 1M HCl and extracted three times with ether. The combined ether extracts were back extracted with 100 ml 1 M aqueous NaOH, and the aqueous phase was then extracted with hexane. The aqueous phase was then acidified with 2 M HCl and extracted twice with ether. The combined ether extracts were washed with brine, dried, and concentrated, yielding (5.1 g, 60%) of the crude acid, which was recrystallized from hexane at −20°C, yielding 3.4 g of fluffy white crystals.

2-methyltetradecanol: A dry flask under Ar was charged with 20 ml dry THF, cooled in an ice bath, and LiAlH_4_ (0.46 g, 12 mmol) was added in two portions. A solution of 2-methyltetradecanoic acid (2.42 g, 10 mmol) in 5 ml THF was then added by syringe pump over 30 min, and the resulting grey slurry was warmed to room temperature and stirred overnight. The mixture was then cooled to 0°C and quenched by sequential dropwise addition of water (0.52 ml), 20% aqueous NaOH (0.39 ml), and water (1.82 ml). The resulting slurry was warmed to room temperature, stirred 30 min, and filtered with suction, rinsing with ether. The filtrate was dried and concentrated, yielding 2-methyltetradecanol (2.18 g, 96%, 99% pure by GC), which was used in the next step without further purification

2-methyltetradecyl iodide: Iodine (3.05 g, 12 mmol) was dissolved in 75 ml CH_2_Cl_2_, and the solution was cooled to ∼-10°C in an ice-salt bath. Triphenylphosphine (3.15 g, 12 mmol) was added in one portion, and the mixture was stirred 1 h. *N*-methylimidazole (0.98 g, 12 mmol) was then added, the mixture was stirred 20 min, and 2-methyltetradecanol (2.12 g, 9.3 mmol) was added in one portion. The mixture was stirred 2 h at ∼-10°C, then quenched with half-saturated aqueous sodium thiosulfate. The CH_2_Cl_2_ was removed by rotary evaporation, and the residue was extracted with hexane. The hexane extract was washed with saturated sodium thiosulphate and brine, dried, and concentrated. The residue was purified by vacuum flash chromatography, eluting with hexane, yielding the desired iodide (2.24 g, 68%, >99% pure by GC).

4-Methylhexadecyne: A dry flask under Ar was charged with 15 ml dry dimethylsulfoxide (DMSO) and lithium acetylide-ethylenediamine complex (1.38 g, 15 mmol), followed by dropwise addition of 2-methyltetradecyl iodide (2.24 g, 6.3 mmol) in 5 ml DMSO over 30 min with a syringe pump. The mixture was stirred for 30 min at room temperature, at which point all the starting material had been consumed. The mixture was quenched by addition of 150 ml ice-water, then extracted with hexane. The hexane solution was washed with 1 M HCl and brine, then dried and concentrated. The residue (1.4 g) was purified by vacuum flash chromatography, eluting with hexane, yielding the desired alkyne (360 mg) contaminated with ∼7% of 2-methyltetradecene from elimination of HI instead of nucleophilic substitution. The alkyne was used in a subsequent step without further purification.

2-Methylpentadecanol: 2-Methylpentadecanol as prepared in 44% yield by alkylation of propanoic acid with tridecylbromide followed by reduction with LiAlH_4_, as described above for the preparation of the homologous 2-methyltetradecanol. The alcohol (1.79 g, 7 mmol) was then converted to the corresponding triflate by treatment with triflic anhydride and pyridine, as described above in the synthesis of 2-methylpentacosane. The crude triflate was taken up in dry ether and used immediately as described below.

7-methyleicos-4-yn-1-ol: A dry flask under Ar was charged with 4-pentyn-1-ol *t*-butyldimethylsilyl ether (1.98 g, 10 mmol), ∼50 mg triphenylmethane indicator, and 25 ml dry THF. The solution was cooled in an ice bath, and BuLi (2.5 M in hexane) was added until the mixture turned pink, followed by addition of a further 4 ml (10 mmol) of BuLi. The mixture was stirred 20 min, then the ethereal solution of 2-methylpentadecyl triflate prepared above was added by syringe pump over 1 h. The mixture was warmed to room temperature overnight, then quenched with saturated aq. NH_4_Cl, and extracted with hexane. The hexane solution was washed with brine, dried, and concentrated. The residue was taken up in 40 ml of a 9:1 mixture of THF and 1 M HCl and stirred at room temperature for 6 h. The mixture was then diluted with water and extracted with hexane. The hexane solution was washed with brine, dried, and concentrated, and the residue was purified by vacuum flash chromatography, eluting first with hexane, then with 17.5% EtOAc in hexane, yielding 7-methyleicos-4-yn-1-ol (0.87 g, 40%, >98% pure by GC). 13,23-Dimethylhexatriacontane: 7-Methyleicos-4-yn-1-ol (0.5 g, 1.62 mmol) was converted to the corresponding triflate as described above and used immediately as described below.

A dry flask under Ar was charged with 4-methylhexadecyne (0.35 g, 1.5 mmol), ∼ 10 mg triphenylmethane indicator, and 10 ml dry THF. The solution was cooled in an ice bath, and BuLi (2.5 M in hexanes) was added dropwise until the solution turned pink, followed by dropwise addition of a further 0.6 ml (1.5 mmol) BuLi. The solution was stirred 20 min, then the ethereal solution of 7-methyleicos-4-yn-1-yl triflate was added by syringe pump over 30 min. When the addition was complete, the mixture was warmed to room temperature and stirred overnight, then quenched by addition of saturated aq. NH_4_Cl, and extracted with hexane. After washing with brine, drying, and concentration, the residue was purified by vacuum flash chromatography, eluting with hexane. The purified product was then taken up in 15 ml hexane, 10% Pd on charcoal catalyst (50 mg) was added, the mixture was flushed with Ar, then H_2_, and then stirred under H_2_ (balloon of H_2_) for 2 h. The mixture was then filtered through celite, and recrystallized from hot acetone, yielding 13,23-dimethylhexatriacontane (110 mg) as white crystals. The synthesis scheme and the EI mass spectrum are shown below.

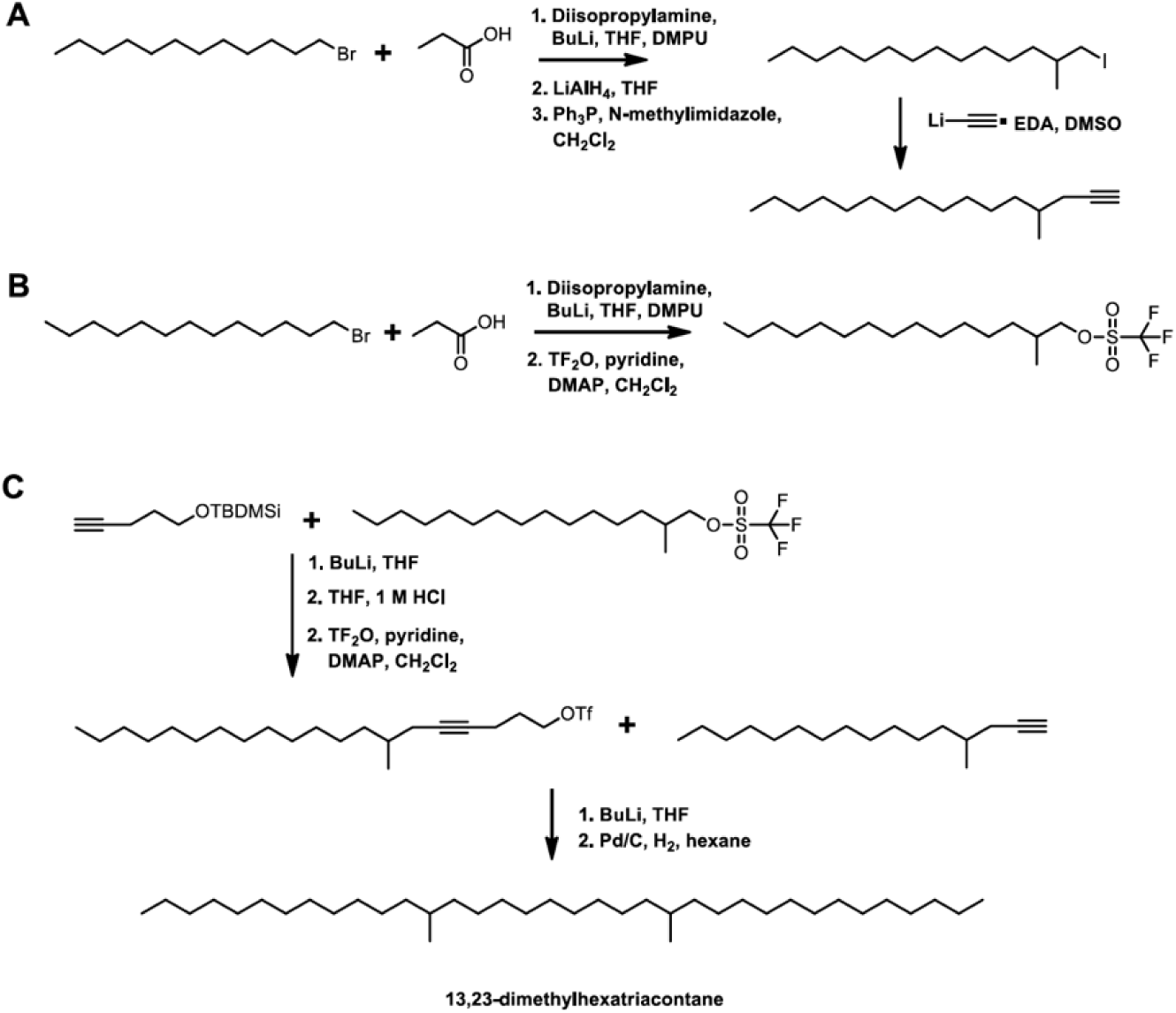

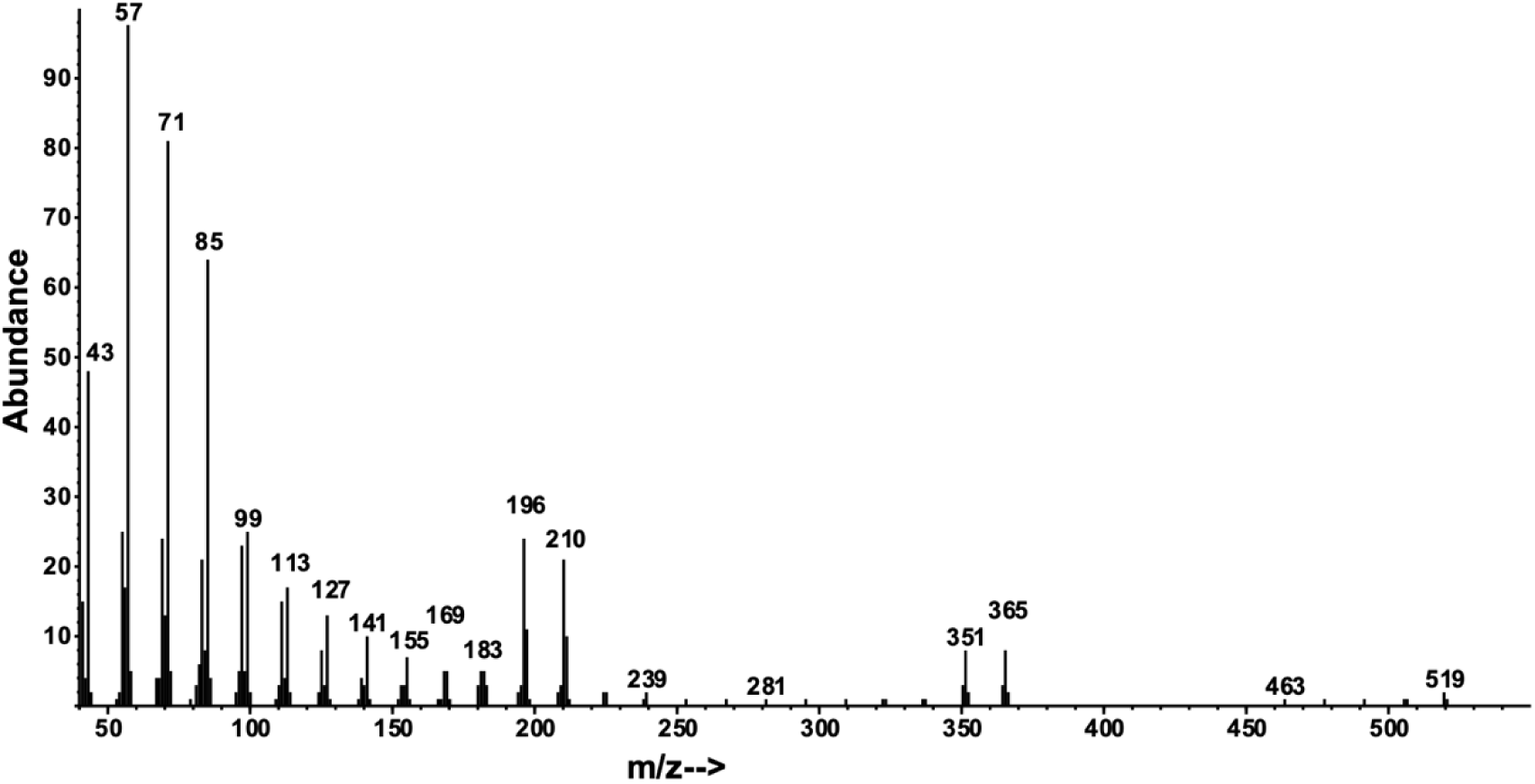

**Dataset S1.** Relative abundance of CHC compounds in hexane extracts from adult ants in the foraging phase, pupae and larvae.

**Dataset S2**. Relative abundance of CHC compounds in hexane extracts from adult ants in the foraging phase and adults in the reproductive phase.

**Dataset S3.** Relative abundance of CHC compounds in hexane extracts from adult ants from different clonal lines and in hexane extracts from nest substrates of clonal line B colonies.

**Video S1.** Representative video of young ants in an arena in which a CHC extract was applied in the shape of the letter “P.”

**Video S2.** Behavioral assay showing a clonal line B ant exploring an arena containing patches treated with CHC extracts from clonal lines A and B.

